# Strain-specific structural variant landscapes shape mutation retention following mutagenesis in *Caenorhabditis elegans*

**DOI:** 10.1101/2025.04.16.649179

**Authors:** R. Kapila, S. Saber, R.K. Verma, G. Blanco, V.K. Eggers, J.L. Fierst

**Author notes:** **<u>Author contributions</u>** J.L.F. secured funding. R.K. and J.L.F. designed the experiments. R.K., with assistance from S.S., performed the fitness assays, and, with assistance from G.B., conducted the male frequency and outcrossing rate assays. V.K.E. carried out the transposable element analysis. R.K. performed the genomic analysis with guidance from J.L.F. R.K.V. developed the Python code for worm counting. J.L.F. developed the theoretical model and conducted the population simulations. R.K. and J.L.F. co-wrote the first draft. R.K. and J.F.K. prepared the final manuscript with contributions from V.K.E. **<u>Competing Interests:</u>** The authors declare no competing financial or non-financial interests.

## Abstract

Classical mutational theories centered on single nucleotide polymorphisms suggest that outcrossing enhances the purging of deleterious mutations by promoting recombination. However, larger structural variants, such as insertions, deletions, and inversions, can suppress recombination and create linkage blocks. Using experimental evolution and whole-genome long- and short-read sequencing, we characterized structural and nucleotide mutation landscapes in three *Caenorhabditis elegans* strains following repeated mutagen exposure and recovery. We found substantial strain-specific differences in structural variant accumulation and mutation retention. The strain with the highest outcrossing propensity exhibited the greatest structural variant burden and a higher fraction of single nucleotide polymorphisms within structural variant intervals. Consistent with this pattern, our population genetic simulations showed that structural variants can persist more readily under higher outcrossing rates. Together, these results indicate that structural variant architecture may influence mutation retention dynamics and highlight strain-specific constraints on purging following mutagenesis in *C. elegans*.

## Introduction

The accumulation of deleterious mutations increases genetic load and can reduce population fitness, potentially leading to long-term decline or extinction ^1,2^. Mutations are broadly categorized into two types: single nucleotide polymorphisms (SNPs) and structural variants (SVs). SNPs are single base pair changes, resulting in localized effects on amino acids or regulatory elements^3^. In contrast, SVs including deletions, duplications, inversions, and translocations, affect larger genomic regions and can alter multiple genes or regulatory networks^4^.

While mutation purging operates on both SNPs and SVs, the dynamics may differ because the majority of SNPs may impact simpler genomic interactions^5^. In contrast, SVs can span larger genomic regions and may suppress recombination, potentially complicating removal^6^. Evolutionary theory has traditionally viewed the spectrum of genomic changes under the broad category of “mutation”, and much of the research on mutation purging has disproportionately focused on SNPs due to their easier detection ^7^. However, the advent of large-scale genome sequencing has revealed the ubiquity of SVs. Despite the foundational role of mutation purging in evolutionary theory^8–10^, how this process operates on SVs remains poorly understood. Addressing this gap is essential for refining theoretical models and understanding how mutation size and severity influence evolutionary dynamics.

Mating systems influence mutation purging through genetic diversity, recombination, and expression patterns of deleterious mutations^11,12^. Population genetics theory predicts that outcrossing populations, through recombination, create novel genetic combinations able to purge deleterious variation^13^. However, outcrossing can also potentially dilute the effects of deleterious mutations and slow purging^14^. In contrast, reduced recombination and genetic diversity in self-fertilizing populations may allow mildly deleterious mutations to accumulate through genetic drift^15^. At the same time, increased homozygosity under selfing can expose harmful recessive mutations to selection and enabling more effective purging^15^. These processes may differ for mutations of different physical sizes asSVs can suppress recombination creating linkage blocks limiting the removal of deleterious variation^16,17^. Consequently, the interaction between mating system, recombination suppression by SVs, and mutation purging remains poorly understood. This complexity is further influenced by transposable elements (TEs), a dynamic source of genomic variation ^18,19^. TEs can generate SVs through transposition or ectopic recombination and may respond to mutagenic or environmental stress^20^. Mating systems may shape TE dynamics: outcrossing can facilitate TE spread but also introduce recombination that helps remove them, whereas selfing can promote TE accumulation through reduced recombination or enhance TE elimination through increased homozygosity ^21,22^. Despite their role in genome evolution, how TEs contribute to mutational landscapes following acute mutagenic stress is also poorly understood.

In this study we use experimental evolution to examine whether strains differing in outcrossing propensity differ in post-mutagenesis mutation retention, and whether these patterns vary with mutation size and genomic context. Specifically, we ask whether strains differing in male frequency and outcrossing rates exhibit distinct patterns of SV accumulation, SNP localization, and mutation retention following repeated mutagen exposure and recovery. By comparing three genetically distinct isolates (N2, AB1, and CB4856), we evaluate whether mutation size and genomic context contribute to strain-specific differences in the persistence of induced mutations. *C. elegans* are androdiecious with populations consisting of both males and self-fertile hermaphrodites. The frequency of males and outcrossing varies drastically across different strains ^23–25^ making *C. elegans* an excellent system to test these ideas. Previous experimental evolution studies in *C. elegans* show that increased mutational input can transiently elevate male frequency and outcrossing, yet does not consistently enhance purging of deleterious variation, with outcomes strongly dependent on genetic background ^26–29^. Comparative genomic analyses across *Caenorhabditis* species further demonstrate that mating system differences shape genome-wide patterns of genetic variation, with self-fertilizing species such as *C. elegans* typically exhibiting reduced nucleotide diversity and stronger linkage among mutations than outcrossing relatives such as *C. remanei* ^30,31^. Together, these studies indicate that reproductive mode can strongly influence mutation dynamics across genomes. However, how these processes affect the retention of different mutation classes—particularly structural variants—following acute mutagenic stress remains largely unexplored.

To test whether strain-specific differences in mating behavior and genomic background influence mutation retention following mutagenesis, we hypothesized that strains differing in outcrossing propensity would exhibit distinct patterns of SNP and SV retention after recovery. We subjected *C. elegans* strains to two mutagens, EMS to induce single nucleotide mutations and formaldehyde to introduce complex lesions in the genome^32^. Populations were mutagenized for five consecutive generations and allowed to recover for three generations. We then tracked relative fitness, changes in male frequency, and outcrossing rates to quantify phenotypic response and investigate patterns of mutation retention after mutagen exposure and recovery. We employed both Pacific Biosystems HIFI long-read and Illumina short-read sequencing to quantify the size, location, and types of mutations that persist after recovery.

## Results

### Strain-specific phenotypic and genomic responses to mutagenesis

To evaluate whether variation in outcrossing propensity is associated with differences in mutation retention following mutagenesis, we subjected three strains of *C. elegans* (N2 (Bristol), AB1, and CB4856 (Hawaiian)), to five generations of mutagenesis followed by three generations of recovery, with four replicate populations for each experimental treatment (12 in total).

### Male frequency was differentially affected by mutagenesis

Male frequency determines the opportunity for outcrossing in *C. elegans* ^24,25^ and changes in male production indicate how mating system dynamics respond to mutagenic stress. Non-mutagenized CB4856 populations had the highest male frequencies (Figure 1A), significantly greater than either AB1 (*p* < 0.0001) or N2 (*p* < 0.01). CB4856 male frequency was reduced after mutagenesis with EMS (*p* = 0.03; Figure 1A) and formaldehyde, although this comparison was not statistically significant (*p* = 0.11). N2 male frequency did not change in response to either mutagen (*p* = 1.00 for both). Similarly, AB1 male frequencies did not differ across populations or treatments (all *p*-values were > 0.05; Supplementary Table 1; Figure 1A).

**Figure 1.**
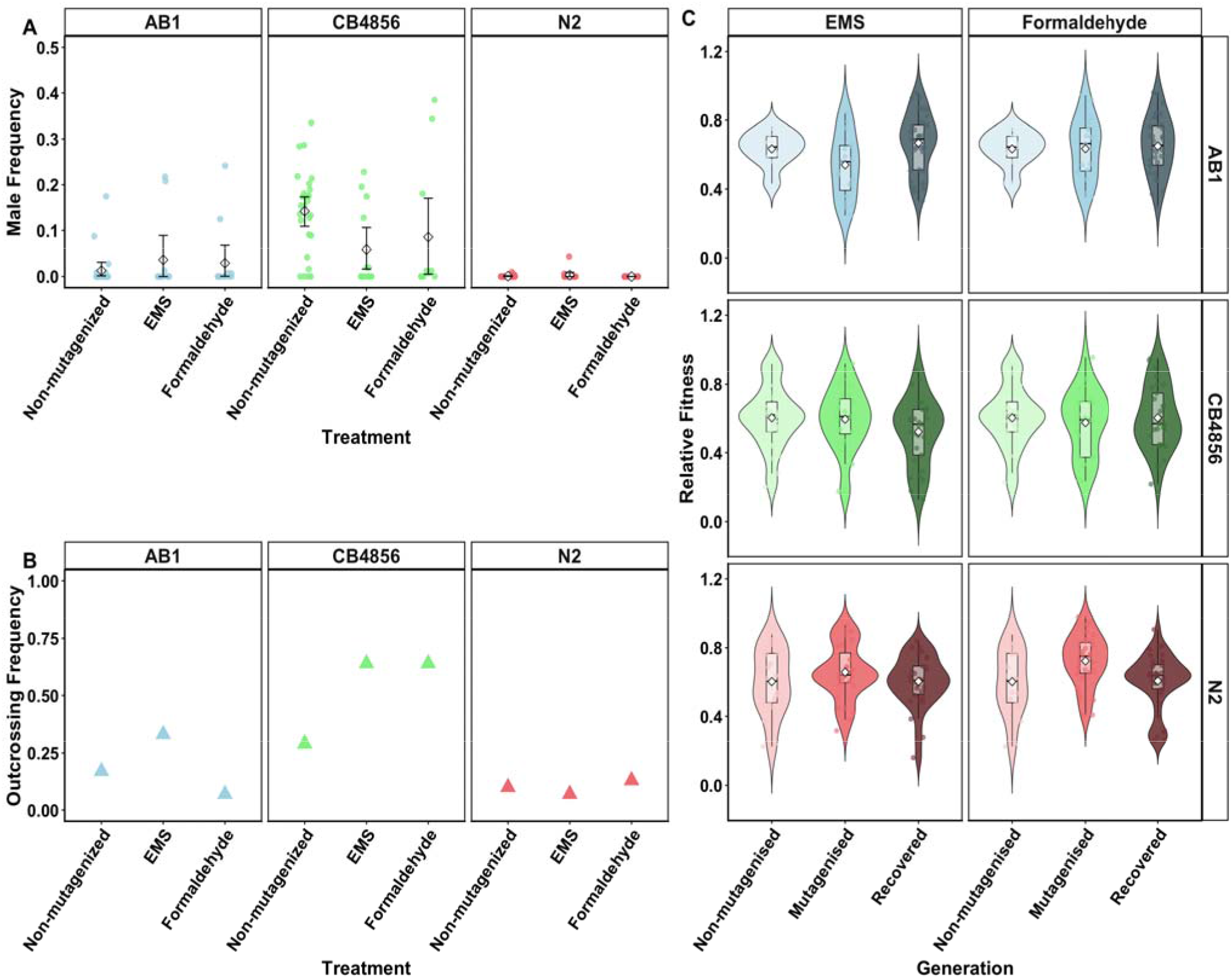
Male frequency, outcrossing behaviour, and relative fitness reveal strain-specific responses to mutagenic stress: (A) Male frequency across strains and treatments. Colored points represent individual assay plates, while open diamonds represent the mean male frequency for each strain–treatment combination. Error bars indicate ±1 standard error (SE) of the mean across plates. Non-mutagenized CB4856 populations exhibited significantly higher male frequencies than AB1 and N2. Mutagenesis had strain-specific effects, reducing male frequency in CB4856 but producing no significant change in AB1 or N2. (B) Outcrossing frequency across strains and treatments. Triangles represent the proportion of assay plates classified as outcrossing for each strain–treatment combination. CB4856 exhibited the highest outcrossing frequencies across conditions, N2 the lowest, and AB1 intermediate levels. (C) Relative fitness distributions across strains and generations. Violin plots show the distribution of plate-level relative fitness values, with boxplots indicating median and interquartile ranges. Colors represent strains and shading indicates generation: non-mutagenized (light), mutagenized (medium), and recovered (dark). Across all strains, relative fitness returned to levels statistically indistinguishable from non-mutagenized baselines following recovery.

### Outcrossing frequencies varied between C. elegans strains

To determine whether mutagenesis altered mating behavior among strains with different male frequencies, we quantified outcrossing frequency using paired mating assays. Cross-fertilization in *C. elegans* produces approximately 50% male progeny while, selfing hermaphrodites produce few or no males. We used male frequency as a proxy for determinig whether outcrossing occurred^33^. In each assay, a single male was paired with a single hermaphrodite, and lifetime male production recorded. Each plate was classified “outcrossing” or “selfing-dominated” using strain-specific male-frequency thresholds established a priori. N2 and AB1 plates with male frequencies >5% were classified as outcrossing. CB4856 exhibited elevated baseline male frequencies and plates were classified as “outcrossing” when male frequency ≥25%. Outcrossing frequency was calculated as the proportion of plates meeting these criteria within each strain and treatment. Thus, outcrossing frequency reflects the probability of successful mating rather than mating efficiency. CB4856 populations had the highest outcrossing frequencies under all conditions, (Figure 1B). In contrast, N2 populations displayed the lowest outcrossing frequencies. AB1 populations were intermediate to these.

### Relative fitness did not differ significantly from parental levels following recovery

Relative fitness, defined here as competitive performance against a GFP-marked reference strain, quantified from relative abundances after ∼8 days of co-culture. Across all strains, recovered populations were statistically indistinguishable from their non-mutagenized ancestors indicating convergence toward baseline fitness (Figure 1C; Supplementary Table 2). CB4856 relative fitness did not differ significantly between non-mutagenized, mutagenized, or recovered populations under either mutagen (all pairwise comparisons p ≥ 0.18). AB1 showed no significant fitnesss differences following either formaldehyde (p = 0.94) or EMS exposure (p = 0.76), although recovered EMS-mutagenized AB1 populations exhibited a modest fitness increase relative to during mutagenesis (p = 0.022). N2 populations exhibited a transient response to formaldehyde exposure, with relative fitness significantly higher immediately after mutagenesis compared to non-mutagenized controls (p = 0.026), an effect not observed under EMS treatment (p = 0.46). After recovery, N2 fitness was statistically indistinguishable from the parental baseline (Recovered vs. Non-mutagenized, p ≥ 0.99).

### Numerous SVs and SNPs were retained after recovery from mutagenesis

Despite rapid fitness recovery, populations harboured numerous *de novo* mutations. CB4856 exhibited significantly higher susceptibility to mutagenesis compared to AB1. When mutagenized with formaldehyde, CB4856 populations showed a 1.12-fold increase in *de novo* SNPs compared with AB1 (t=5.55, *p*=0.0014) and comparable *de novo* SNP counts to N2 (t < 0.01, *p*= 0.996). In contrast, with EMS as the mutagen, CB4856 showed a significantly higher *de novo* SNP count when compared with AB1 (t = 6.79, p < 0.01) and a marginally higher *de novo* SNP count when compared with N2 (t = 2.45, p = 0.05; Figure 2A).

**Figure 2.**
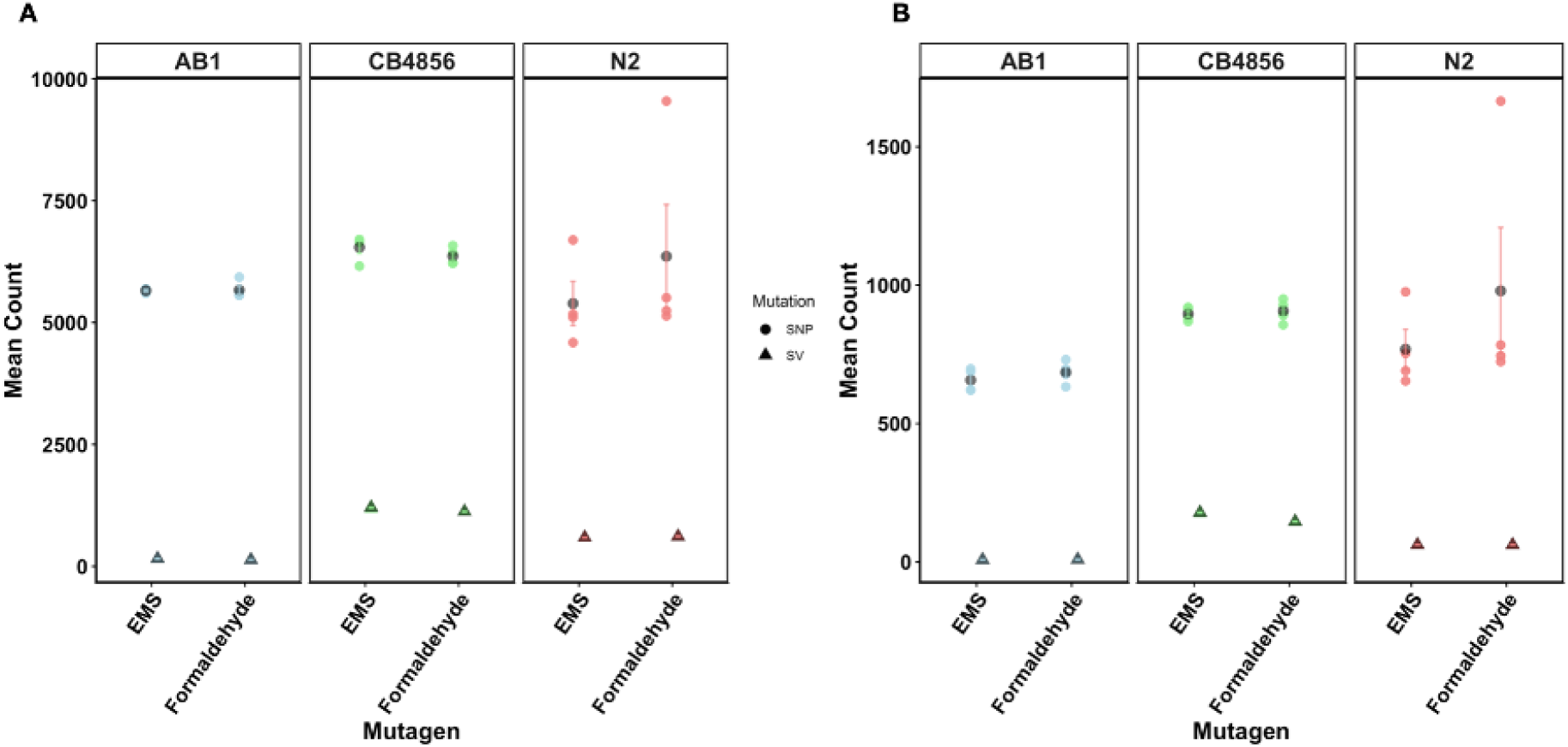
Comparison of de novo SNPs and SVs mutations and their localization in exons across strains (A) Mean counts of de novo SNPs (Circles) and SVs (Triangles) induced by EMS and Formaldehyde across AB1 (Green), CB4856 (Red), and N2 (Blue). De novo SNPs are significantly more abundant than de novo SVs under both mutagen treatments. CB4856 has the maximum number of de novo SNPs and de novo SVs. (B) Mean number of de novo SNPs and de novo SVs localized in exons for the same strains and mutagen treatments, with de novo SNPs again being more prevalent than de novo SVs and CB4856 having more SVs in exons than the other two strains and more SNPs in exons than AB1. Compared to N2, CB4856 had more SNPs in exon than N2 when treated with EMS and comparable SNPs in exon when treated with Formaldehyde.

Due to marked heteroscedasticity in SV counts we performed Mann–Whitney U tests to statistically evaluate *de novo* SV counts. CB4856 populations had significantly higher *de novo* SV counts (Figure 2A) when compared with AB1 (W=16, *p* = 0.03 for both mutagens) and N2 populations (W=16, *p* = 0.03 for both mutagens).

Across strains and treatments, most of the detected SVs were intrachromosomal insertions, deletions, duplications and inversions. Interchromosomal translocations were just ∼0.7–4.3% of all identified SVs (Supplementary Table 3). Thus, most mutagen-induced SVs reflect within-chromosome rearrangements rather than widespread between-chromosome exchanges.

### CB4856 harbored high numbers of SNPs and SVs in exons

To evaluate whether mutation retention differed across functional genomic regions, we examined the distribution of SNPs and SVs relative to annotated exons. CB4856 populations had the highest number of mutations in coding regions (Figure 2B) and the highest number of SNPs located within SVs (Figure 3A). CB4856 had 2.20 times more exonic SNPs than AB1 in formaldehyde-mutagenized population (W=0, *p*= 0.03) and 1.12 times more than N2 (W=12, *p*=0.34). CB4856 had 2.26 times more exonic SNPs compared to AB1 in EMS mutagenized population (W=0, *p*=0.03) and 1.38 times more exonic SNPs compared to N2 (W=16, *p*=0.03) (Figure 2B).

**Figure 3.**
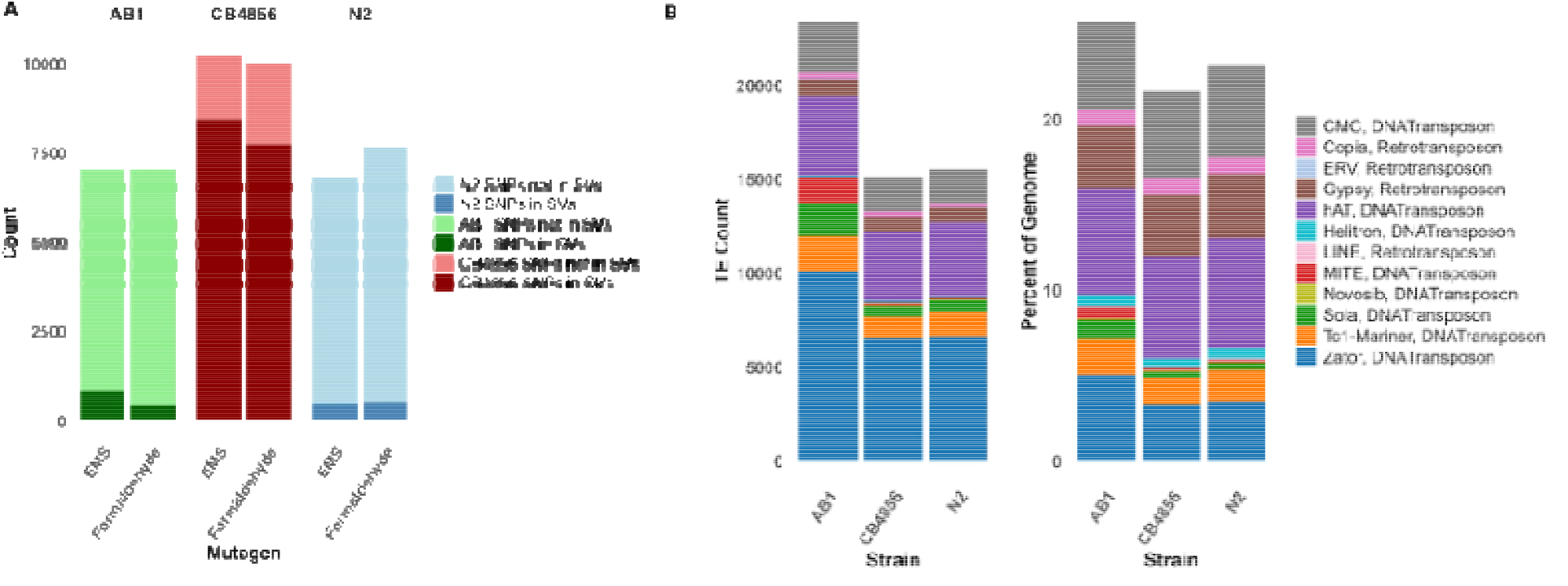
Mutational and TE landscape across strains and mutagens **(A):** Number of SNPs, SVs and SNPs in SVs for AB1, CB4856 and N2; **(B):** Classification and count of overall TE families contributing to transposition events across strain-treatment combinations. DNA transposons such as Zator and Tc1-Mariner were among the most active elements, with CB4856 showing the broadest TE activity profile.

Exons were similarly affected by SVs. AB1 populations had fewest exons containing SVs, whereas CB4856 populations had the most number. AB1 formaldehyde populations had 16.25-fold fewer exonic SVs than CB4856 (W = 0, p = 0.02) and 6.92-fold fewer than N2 (W = 16, p = 0.03). CB4856 EMS populations had 22.31-fold more exonic SVs compared to AB1 (W = 0, p = 0.02), and N2 had 7.81-fold more when compared with AB1 (W = 16, p = 0.03; Figure 2B).

CB4856 had the highest number of SNPs located within SVs interval (77.13% for formaldehyde and 82.46% for EMS treatment; Figure 3A). SVs in AB1 populations contained 6.36% (formaldehyde) and 12.01% (EMS) of *de novo* SNPs. N2 populations had similarly low numbers of SNPs contained in SVs (EMS 7.12%; formaldehyde 6.80%).

### Non-random enrichment of SNPs within structural variant regions

SNP counts within SV intervals were significantly greater than random expectation (all p < 0.001; Supplementary Table 4). Enrichment ranged from 2.6–3.0× in AB1, 2.6–4.9× in N2, and ∼1.9× in CB4856 (Supplementary Table 4). CB4856 exhibited substantially greater SV genome coverage (∼40– 43%) (Figure 4) with SNPs significantly enriched within SV regions relative to genomic expectation.

**Figure 4.**
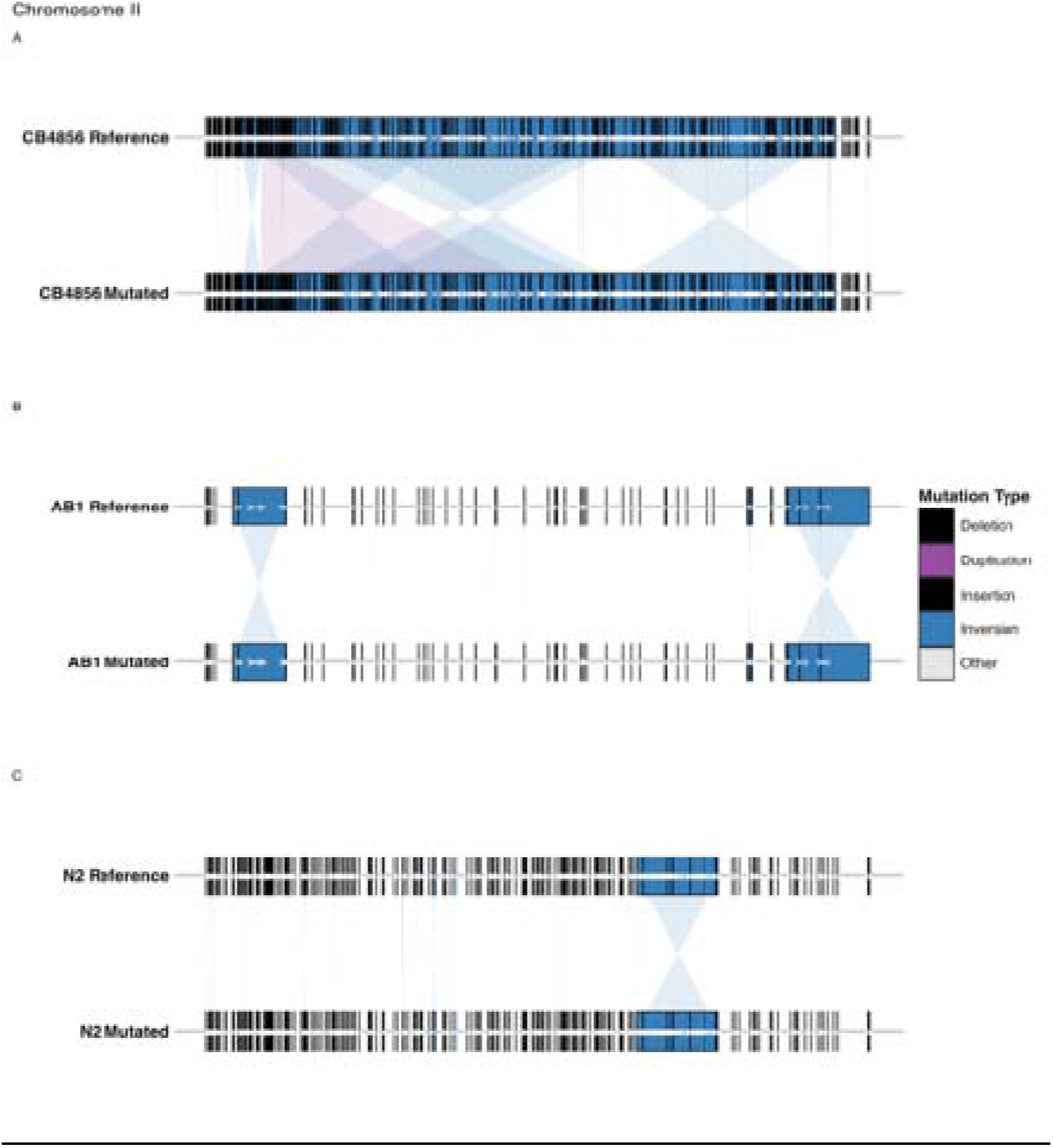
Structural variation patterns across chromosome II following EMS mutagenesis. Comparison of structural variants detected in reference and mutagenized populations of CB4856, AB1, and N2. Black tick marks indicate individual structural variant breakpoints, while colored blocks denote larger structural rearrangements classified by mutation type (deletion, duplication, insertion, inversion, or other). Shaded connectors indicate large inversion or rearrangement events between reference and mutagenized genomes. CB4856 populations exhibit a greater density and span of structural variants relative to AB1 and N2.

### Breakpoint microhomology differed among strains but did not explain SV burden

To test whether strain-specific differences in SV abundance reflect differences in DNA repair signatures, we examined breakpoint microhomology across intrachromosomal SVs. Non-Homologous End Joining (NHEJ) results in 0–2 bp mutational ‘scars’ while Microhomology-Mediated End Joining (MMEJ) leaves larger 3–10 bp signatures. Homology-mediated repair such as Single-Strand Annealing (SSA) results in ∼11-50 bp mutations while mutations >50bp indicate different mechanisms of Homology-Mediated Repair (HMR). Accordingly, we classified breakpoints into bins of 0 bp (NHEJ) 1–2 bp (NHEJ/MMEJ), 3–10 bp (MMEJ), 11–50 bp (SSA), and >50 bp (HMR) (Supplementary Table 5). NHEJ and NHEJ/MMEJ occurred at comparable frequencies across strains. MMEJ and SSA tracts were enriched in AB1 while CB4856 and N2 were enriched for larger HMR signatures. N2 had the highest enrichment of extended HMR mutations despite having an intermediate overall SV burden, indicating that breakpoint architecture alone does not explain strain differences in SV abundance.

### Active transposable elements were responsible for a small portion of SV mutations

Because both mutagenesis and outcrossing can influence transposable element (TE) activity, we evaluated whether TE mobilization contributed to SVs in mutagenized populations using the TransposonUltimate pipeline ^34^.

Total repeat content comprised 23.21%, 25.74%, and 21.70% of the N2, AB1, and CB4856 reference genomes, respectively. Class II DNA transposons represented ∼94% of annotated TEs in each genome (Figure 3B). Zator elements were the most abundant, accounting for ∼46% of DNA TEs. AB1 contained a smaller proportion of hAT elements but a higher proportion of Sola and MITE elements relative to N2 and CB4856. Tc1-Mariner (9%), CMC (13%), Helitrons (0.5%), and Novosib (0.3%) elements occurred at similar frequencies across genomes. As expected for *C. elegans*, TE density followed the characteristic chromosome arm–center pattern, with higher TE abundance on chromosome arms (Figure 5).

**Figure 5.**
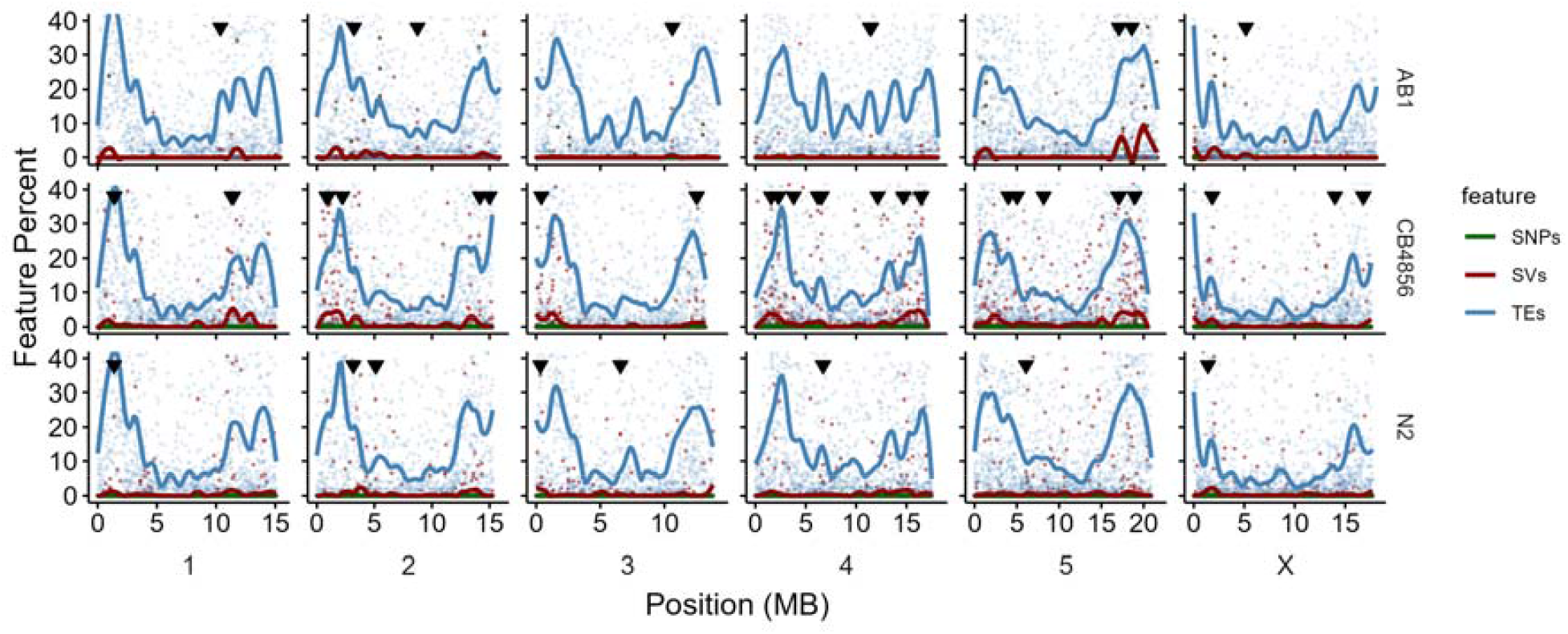
Genomic distribution of SNPs, structural variants (SVs), and transposable elements (TEs) across the genomes of three Caenorhabditis elegans strains following EMS mutagenesis. Columns represent the six chromosomes, and rows correspond to the three strains (AB1, CB4856, and N2). Points show the percentage of 10-kb genomic windows containing each feature across genomic position. Smoothed lines represent locally estimated scatterplot smoothing (LOESS) fits for transposable elements (blue), structural variants (red), and SNPs (green). Black downward-facing triangles indicate positions of predicted active transposable elements. Features exceeding 10 kb in length were excluded from the visualization; details of these larger structural variants are provided in Supplementary Table 9.

To assess TE mobilization after mutagenesis, we intersected reference TE annotations with de novo SV calls using the TEdeTEct pipeline implemented in TransposonUltimate. Only a small fraction of annotated TEs showed evidence of transposition: 0.07–0.08% in N2, 0.04–0.05% in AB1, and 0.24–0.27% in CB4856. Consequently, TE-associated events accounted for <15% of all de novo SVs. Despite having the lowest genomic TE proportion, CB4856 exhibited ∼4× more transposition events than N2 or AB1 (Supplementary Table 6).

TE activity was broadly similar between EMS and formaldehyde treatments. In N2, 6 of 21 (29%) transposition events occurred under both mutagens. CB4856 and AB1 showed 12 of 67 (18%) and 1 of 22 (4.5%) shared events between treatments, respectively, and no identical transposition events were observed across strains. Zator elements accounted for ∼55% of all detected transposition events (Figure 3B). Together, these results indicate that TE activity contributes to the formation of de novo SVs but, it accounts for a minority of the SVs observed following mutagenesis.

### Mutagenesis induced large SV mutations

Although most SVs were small (<10^4^ bp), both formaldehyde and EMS generated large-scale mutations affecting substantial portions of the genome (Supplementary Figure 1). Across all strains and treatments, SVs larger than 10 kb accounted for only ∼1.4% of detected events (Supplementary Table 7), yet these included several megabase-scale inversions capable of reshaping large genomic regions (Figure 4).

In the N2 formaldehyde-treated population, insertion added 471,238 nucleotides (∼0.48% of the 100.8 Mb *C. elegans* genome). In comparison, 74,044 nucleotides (0.06%) were deleted, 44,318 nucleotides (0.04%) duplicated, and 1,289,368 nucleotides (1.29%) affected by inversions. Mutations in the EMS-mutagenized N2 populations were of similar scale (Supplementary Table 8). Large inversion events were also detected in N2 populations, including a 1.09 Mb inversion on chromosome IV in formaldehyde-treated populations and a 2.49 Mb inversion on chromosome × in EMS-treated populations (Supplementary Table 9).

AB1 populations had the smallest number of nucleotides affected by SV mutations. In formaldehyde-treated AB1, 35,468 nucleotides were inserted and 46,363 deleted. The notable exception was inversion, which spanned 2,539,375 nucleotides in formaldehyde-treated and 4,181,603 nucleotides in EMS-treated populations. Some transposition events were shared between mutagen treatments. For example, a 2.54 Mb inversion on the right arm of chromosome 5 occurred in both AB1 Formaldehyde and EMS-mutagenized populations. AB1 EMS-mutagenized populations also experienced a 1.56 Mb inversion on the left arm of chromosome 4 (Supplementary Table 9).

CB4856 populations harbored substantially more DNA affected by SVs (Figure 4; Supplementary Table 3). Roughly 1 Mb of new DNA was inserted under both treatments (Supplementary Table 3). Compared to AB1, CB4856 had 14.11-fold more insertions under formaldehyde and 15.18-fold more under EMS. Relative to N2, it had 1.50-fold and 1.66-fold more insertions under formaldehyde and EMS, respectively. CB4856 also contained large amounts of duplicated DNA, totaling 1.7 Mb under formaldehyde and 21.7 Mb under EMS, largely driven by a 12.31 Mb duplication on chromosome IV and an 8.82 Mb duplication on chromosome I. Inversions represented the largest class of structural change, spanning 49.7 Mb—nearly half the genome—in formaldehyde-treated CB4856 populations, including 13.91 Mb and 7.1 Mb inversions on chromosome V, an 11.24 Mb inversion on chromosome III, and 10.5 Mb and 6.29 Mb inversions on chromosome II. In EMS-treated CB4856 populations, inversions spanned 32.8 Mb—nearly one-third of the genome—including large inversions on chromosomes II, I, and V (Supplementary Table 9).

When SVs larger than 10 kb were excluded, the remaining mutations followed the characteristic arms–center pattern of the *C. elegans* genome, with more nucleotides affected on chromosome arms than in centers (Figure 5), consistent with the recombination landscape where recombination is elevated on arms and suppressed in central regions. However, the presence of multiple megabase-scale SVs indicates that mutagenesis generates structural changes capable of influencing recombination and mutation linkage across large genomic regions.

### SVs and transposon activity varied across chromosomes

The distribution of SVs varied across chromosomes, strains, and mutagens (Figure 6A). SV mutations were more frequent on Chromosomes IV and V compared with Chromosome X. TE activity did not explain these patterns; for example, TEs were most active on Chromosome IV in CB4856, whereas Chromosome V harbored more SVs (Figure 6B). TE-associated duplications and inversions were less frequent than deletions across all populations. These patterns indicate that TE activity alone does not account for the chromosomal distribution of SVs.

**Figure 6.**
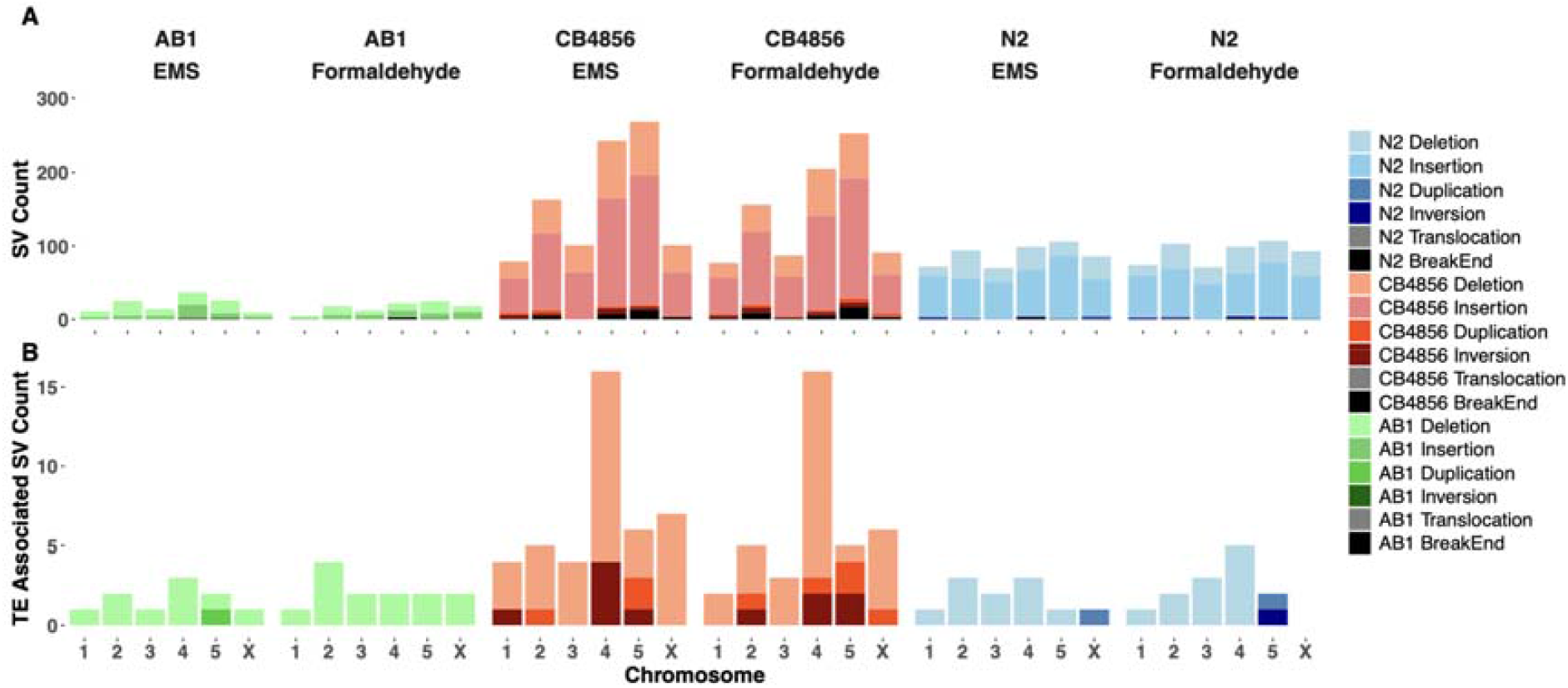
Chromosomal distribution of SVs and TE-associated SVs across strains and mutagens. (A) Stacked bar plots showing the total count and types of SVs including Deletions, Insertions, Duplications, Inversions, Translocations, and Unclassified (NA) across chromosomes 1–5 and X, for each strain (AB1, CB4856, N2) and mutagen (EMS and formaldehyde). (B) TE associated SVs were primarily Deletions.

### Simulation support of experimental results

Although there is a body of work modelling the influence of outcrossing on mutation in populations these models typically assume that mutations are occurring at the level of an individual SNP or broader, abstract ‘allele’ level^35–37^ . To explore whether our experimental results could emerge under different evolutionary scenarios, we developed a population model describing evolution of SVs in the genome. We implemented this model in stochastic simulations using the SLiM 4.3 software^38^ and measured the impact of outcrossing rates on SVs in the population.

Consistent with our experimental observations, the simulations showed that outcrossing populations retained SVs, particularly when outcrossing rates consistently exceeded 30%. In contrast, fully selfing populations rapidly purged SVs. SVs were retained in outcrossing populations when each mutation carried a small deleterious effect on fitness, drawn from a gamma distribution^39–41^ (Supplementary Figure 2). The fitness landscape had a negative curvature or synergistic epistasis (Supplementary Figure 3). Under these conditions, SVs persisted in simulated populations. These results suggest that outcrossing can, under specific conditions, promote the persistence of structural mutations (Figure 7).

**Figure 7.**
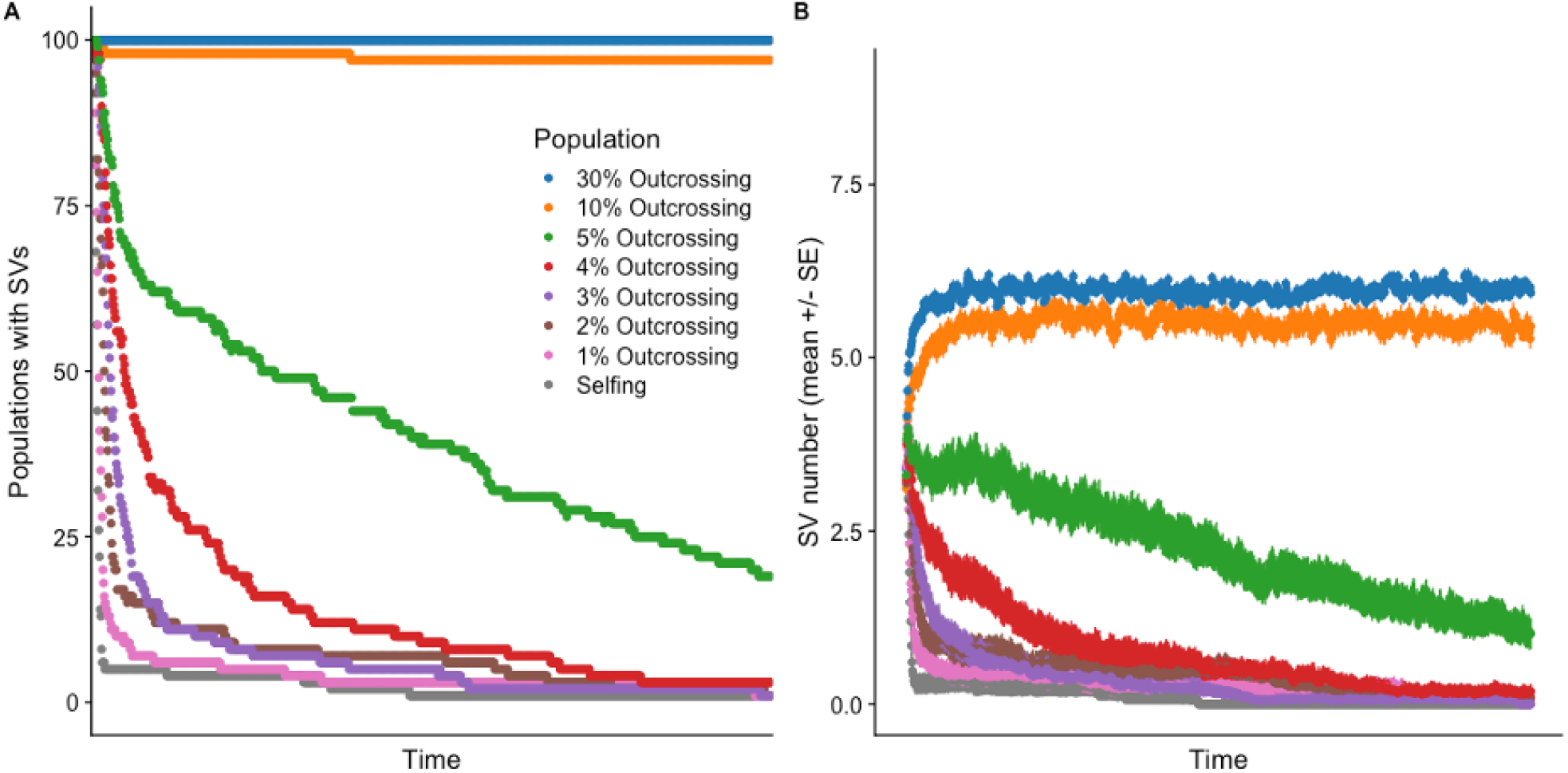
A) Simulated populations evolving with less than 10% of individuals reproducing through outcrossing do not retain SVs. B) The mean number of structural variant mutations is lower in populations with lower quantitative levels of reproduction through outcrossing. Here *K*=1000, =0.001, =0.0006, =0.01, =0.001 and recombination rate =0.0001.

## Discussion

In this study we compared genomic responses to repeated mutagen exposure and recovery in three genetically distinct *C. elegans* isolates. Integrating experimental evolution, whole-genome sequencing, and population genetic simulations revealed strain-specific differences in mutation retention associated with mating behavior and genomic background. CB4856, which exhibited the highest male frequency and outcrossing rate, retained large numbers of mutations despite recovering relative fitness after mutagenesis. Notably, many SNPs occurred within SV intervals and frequently overlapped exonic regions. Although SNP enrichment within SV regions occurred in all strains, the absolute number was far greater in CB4856, consistent with genome wide SVs coverage suggesting that genome architecture influences mutation retention following mutagenesis.

*C. elegans* has been widely used to study mutation accumulation and genomic responses to mutational stress through both mutation-accumulation experiments and analyses of naturally occurring variation ^42,43^. Experimental evolution studies show that increased mutational input can elevate male frequency and outcrossing rates, likely as short-term responses to mutational load ^26,29^. However, elevated outcrossing does not consistently result in sustained purging of deleterious mutations, and natural variation among isolates highlights strong genetic background effects ^26,28,29^. Our results extend these findings by showing that strain-specific genomic architecture can influence genome-wide patterns of mutation retention following mutagenesis.

We explored these factors in our population simulations and found that under specific assumptions increased levels of outcrossing would retain SVs. Importantly, the phenotypic effect size of the segregating mutations was very small. In a fully outcrossing simulated population, there was an average of 10 mutations per individual after 100,000 generations, each with an average phenotypic effect size of 0.0035. Larger mutation effect sizes were rapidly purged from outcrossing populations. However, deleterious mutations with small effects on the phenotype and fitness were stably maintained in the simulated outcrossing populations for long periods. Because mating system was explicitly manipulated in the simulations but not in the experimental populations, we interpret the simulations as providing mechanistic support for (rather than direct validation of) the empirical associations observed among strains.

Genomic responses to mutational stress depend in part on the type of DNA lesion and the repair mechanisms that resolve it. Point mutations are typically repaired through efficient pathways like base excision repair (BER) or nucleotide excision repair (NER), while complex DNA lesions generating double-strand breaks must be resolved through pathways such as homologous recombination (HR), classical non-homologous end joining (NHEJ), and microhomology-mediated end joining (MMEJ), which can produce SVs during repair^44^. We expected that EMS should create more SNPs while formaldehyde would create more SVs in the genome. However, we found that 5 consecutive generations of treatment with either chemical had similar mutational effects on the *C. elegans* genome. Breakpoint microhomology analysis revealed that homology patterns differed among strains but did not correspond directly with SV abundance (Supplementary Material 7). In particular, CB4856 exhibited the highest SV burden despite displaying breakpoint signatures broadly similar to those observed in N2. These results suggest that differences in SV abundance are unlikely to arise solely from variation in repair pathway usage and may instead reflect differences in genome architecture.

SV in *Caenorhabditis* is dominated by intrachromosomal rearrangements, whereas interchromosomal translocations are rare^46^. Recent mutation-accumulation studies provide benchmarks for spontaneous SVs in *C. elegans*^*47*^, which occur at low frequency and are dominated by insertions and deletions. Consistent with this pattern, the SVs detected following mutagenesis in our study were overwhelmingly intrachromosomal across all strains and treatments (Supplementary Table 4). Comparisons between the CB4856 and N2 reference genomes have used long-read DNA sequencing to reveal extensive structural divergence at the same time as chromosome-scale synteny is preserved^48,49^, analysis of the non-mutagenized populations mirrored this pattern, with CB4856 harboring substantially greater standing SV variation than N2. The mutagen-induced SVs we observed exhibited similar qualitative properties: across strains and treatments, SVs were primarily insertions and deletions, intrachromosomal rearrangements predominated, and exon overlap occurred at frequencies comparable to those reported for spontaneous structural mutations^47^ (Supplementary Tables 3–10). These similarities suggest that mutagenesis increases the frequency and genomic impact of SVs without generating qualitatively different classes of rearrangements. These strains also differ extensively at the nucleotide level, with hundreds of thousands of SNPs and numerous short insertions and deletions distributed across the genome ^49,50^. Because long-read sequencing was performed at a single post-recovery time point, our data capture the net genomic outcome of recovery from mutagenesis rather than directly measuring purging rates. Nevertheless, all strains recovered relative fitness within three generations, indicating no irreversible fitness decline. Differences in mutational load across strains could, in principle, reflect variation in initial susceptibility to mutagenesis or DNA repair efficiency, in addition to differences in purging dynamics. However, all strains recovered relative fitness within three generations following mutagenesis, indicating that none suffered irreversible fitness impairment under these conditions. Moreover, population genetic simulations that hold repair efficacy constant recapitulated the empirical observation that higher outcrossing rates are sufficient to retain SV that would otherwise be purged in predominantly selfing populations, even in the absence of strain-specific differences in DNA repair.

AB1 exhibited relatively low numbers of SNPs and SVs, limited exonic overlap, and moderate SNP enrichment within SV regions (∼2.5–3× above expectation). Because SVs span a relatively small fraction of the genome and contain comparatively few coding mutations, this configuration may reduce functional disruption. In this context, SNP enrichment likely reflects localized clustering of mutations within a comparatively constrained mutational landscape, potentially buffering AB1 against large-scale functional perturbation following mutagenesis.

In CB4856 populations, SVs spanned approximately 40–43% of the genome and contained the highest absolute number of SNPs embedded within SV regions. Although enrichment relative to genomic expectation (∼1.9×) was lower than in N2, the overall burden of coupled structural and nucleotide variation was substantially greater. Many of these variants intersected coding regions, increasing the likelihood of functional disruption. SVs in CB4856 also spanned a broad size range (10^3^–10^7^ bp; Supplementary Figure 1), generating a genome-wide mutational landscape that may promote genomic plasticity but also increase susceptibility to instability under chemical mutagenesis. Despite this elevated mutational burden, CB4856 populations recovered relative fitness to levels indistinguishable from the ancestral baseline within three generations, indicating that short-term fitness does not fully capture underlying genomic load.

N2 presents a contrasting genomic response. Although SVs span only a small proportion of the genome (∼1–3%), SNP enrichment within SV regions was among the highest observed (∼2.6–4.9× above expectations), indicating strong local clustering of nucleotide variation within structurally altered segments despite limited overall SV coverage. Because the total number and genomic span of SVs remain relatively constrained, the absolute burden of linked structural and nucleotide variation is potentially lower than in CB4856. N2 populations also showed relatively stable male frequencies and rapid fitness recovery, suggesting that SVs were spatially restricted. Interestingly, relative fitness in N2 increased transiently following formaldehyde exposure before returning to ancestral levels after recovery (Fig. 1B). This short-term increase may reflect consequences of long-term laboratory domestication. Unlike the wild isolates AB1 and CB4856, N2 has experienced decades of selection in stable laboratory environments^51^, whereas CB4856 represents a genetically diverged natural isolate^52^. The transient increase in relative fitness observed in N2 following formaldehyde exposure may reflect a compensatory physiological response analogous to hormesis, in which short-term stress induces beneficial physiological effects^53,54^.

Our observation that SNPs are consistently enriched within (SV) regions across strains suggests that nucleotide and structural mutations frequently occur within the same genomic intervals. Such clustering can reduce the effective separation of linked mutations during recombination, particularly when SVs span large genomic regions. These patterns provide a genomic context for previous experimental evolution studies in *C. elegans*, which show that increased mutational input can elevate male frequency and outcrossing but does not consistently enhance purging of deleterious mutations ^26,28,29^. Together, these results suggest that the physical organization of mutations within genomes may influence how efficiently recombination can separate and remove deleterious variants following mutagenic stress.

Transposable elements (TEs) can play a critical role in shaping genomic resilience by introducing genetic variation and driving structural changes in response to stress^55^. Increased outcrossing has been associated with elevated TE activity, likely due to the enhanced recombination rates and reduced efficiency of silencing mechanisms in outcrossing populations^56,57^. Our results show strain-specific differences in TE activity following mutagenesis. CB4856, which had the highest outcrossing frequency, also exhibited the highest transposition rates, consistent with increased genomic instability under mutagenic stress. In contrast, although AB1 had the highest number of annotated transposons, its transposition rates were lower, suggesting fewer active TEs. N2 showed the highest overlap in TE events between mutagens (∼30%), suggesting common TE activations under stress.

Unlike earlier studies focusing on the Tc1-Mariner family of TEs ^58,59^, we found that Zator was the most active and abundant transposon across all three strains and both mutagens. This result is consistent with recent work showing that few TEs remain active in *C. elegans*, most of which belong to the Zator family^49,60^. The transposon landscape also showed the expected arm–center pattern reported in previous studies, consistent with the known recombination landscape of *C. elegans*, in which recombination is elevated on chromosome arms and reduced in chromosome centers^61^. Together, these observations suggest that elevated TE activity in CB4856 may contribute to its increased genomic instability under mutagenic stress. However, because TEs account for only a small fraction of the total SVs in our dataset, TE mobilization is unlikely to explain the full extent of strain differences in mutation retention.

In conclusion, we found strain specific fitness and mutation responses which may reflect larger patterns of evolutionary response. Although the N2 “Bristol” strain, originally collected by Sydney Brenner^51^ has been developed as a model system, its fitness, behavior and genomics differ substantially from other strains of *C. elegans*. N2 is a highly lab adapted strain with little genetic variation, few males and low outcrossing frequencies ^62^. In contrast, CB4856 and AB1 are more recently collected “wild worms” with substantial genomic differences ^50^ and high natural variation in male frequency and outcrossing ^24,28^. Additionally, our analyses indicate that SNPs are significantly enriched within SV regions relative to genomic expectations, suggesting that strain specific genomic landscapes determine the region in which mutations accumulate and remain linked. Such clustering may influence how mutational load is distributed across the genome and how efficiently outcrossing and recombination can separate deleterious variants following mutagenic stress. However, because these comparisons involve distinct genetic backgrounds, the present results do not establish a causal relationship between outcrossing and SV persistence. Forward-time simulations were therefore used to explore how variation in mating system parameters could influence SV persistence under controlled conditions; these are presented as a mechanistic framework consistent with, but not demonstrating, causality. Future studies manipulating mating system within a common genetic background, for example using *him* mutants to increase male frequency or *mab* mutants to prevent mating, together with strain-specific analyses using *fog* or *xol* backgrounds and additional mutagenic treatments, will help clarify how mating dynamics influence mutation retention. Overall, our results suggest that genome architecture and mating system jointly shape how populations respond to mutational stress.

## Supporting information

Supplementary figures tables and protocols

## Acknowledgements

The authors would like to thank the members of the Fierst lab and Jessica Gonzalez, Jason Pienaar and Jesualdo Fuentes-Gonzalez for suggestions and careful feedback. We also thank four anonymous reviewers who provided thorough review and greatly improved the work.

## Data Availability

Scripts, bioinformatic workflows and analyses are available at github.com/RohitKapila. Python code used for image analysis is available at github.com/FierstLab/WormImage. Fitness data are available as Supplemental Materials associated with the manuscript. Sequence reads are available at the NCBI SRA under BioProject PRJNA1201181.

## Methods

### Strains and mutagens

Three *C. elegans* strains with varying male frequencies were chosen: N2 (Bristol), CB4856 (Hawaiian), and AB1. ST2 served as a common competitor in relative fitness assays. All strains were obtained from the Caenorhabditis Genetics Center and maintained frozen until the experiment. Frozen strains were thawed on 75 mm × 13 mm NGM plates seeded with *E. coli* OP50. After one generation (∼3.5 days at 20 °C) for recovery, each strain was divided into nine replicate populations. Prior to the experiment, replicates were synchronized by bleaching. Over five generations, four replicates per strain were mutagenized with formaldehyde, four with EMS, and one remained untreated as a non-mutagenized control. All replicates for a given strain were handled in parallel on the same day using the same batch of plates to minimize intra-strain variability due to plate effects. A single non-mutagenized replicate was included per strain as a reference baseline rather than as a fully replicated treatment. This design choice reflects the extremely low rates of spontaneous SNP and structural variant (SV) accumulation in *C. elegans* (approximately 0.003–0.03 SVs per genome per generation)^42,47^. Over the five-generation duration of this experiment, the expected number of spontaneous de novo SVs is therefore near zero. Moreover, because genomic analyses were performed using pooled long-read sequencing, any spontaneous variants arising in untreated populations would be present at very low frequencies and fall below reliable detection thresholds. Under these conditions, additional untreated replicates would be expected to accumulate negligible mutational differences and provide little additional biological information. Replication was instead prioritized within mutagenized treatments to maximize power and sequencing depth for detecting treatment-specific genomic and fitness consequences. Following the mutagenesis phase, all populations were frozen at −80 °C for downstream analyses.

### Mutagenesis

Four replicates per strain were treated with 1 mM ethyl methanesulfonate (EMS) in M9 solution, while the other four replicates were treated with 0.1% (w/v) formaldehyde solution in M9. These concentrations were chosen because they are non-lethal yet sufficient to induce detectable genomic mutations. At each transfer, entire populations were moved in suspension (300 µL per replicate), preserving large census sizes and minimizing bottlenecks. Based on subsampling counts of the transfer suspension, populations were estimated to contain approximately 3,000–4,000 individuals per replicate Mutagenesis was carried out for five consecutive generations. During this phase, populations were exposed to the mutagen for 4 hours every fourth day before being transferred back to standard NGM plates seeded with *E. coli* OP50. Following the mutagenesis phase, all populations were allowed to recover for three additional generations without mutagen exposure, during which worms were maintained under standard laboratory conditions. We followed the worm mutagenesis protocol described by Brenner et al. (1974) with minor modifications. Every generation (∼4 days), worms were washed from the plates with M9, collected in tubes, and resuspended in their respective mutagen solutions for 4 hours at 20 °C, during which they were gently rocked to ensure uniform exposure. After treatment, worms were transferred to fresh plates with NGM and *E. coli* OP50, ensuring maintenance of replicate identity and avoidance of mixing. Population transfers were performed without developmental synchronization. Generations were overlapping, i.e., individuals at different developmental stages contributed to subsequent generations.Population sizes were not altered at any point during the experiment. During both transfers, from the plates to the tubes and from the tubes back to the plates, all worms were transferred using 300 microliters of liquid. To approximate census size, we counted worms in replicate 10-µL subsamples of the transfer suspension and extrapolated to 300 µL (average ≈ 3,300 worms per transfer). Specifically, M9 buffer was used for the transfer from plates to tubes, and the mutagen was used for the transfer from tubes back to plates. After the mutagen treatment, the liquid was carefully transferred onto the plate, causing parts of the *E. coli* colony to be dislodged. As a result, the worms would initially swim on the plate. We then left the plate open, allowing the liquid to be absorbed within 30-40 minutes, after which the worms resumed crawling on the plate.

### Male frequency assay

To evaluate the impact of five generations of mutational exposure on male frequency, a male frequency assay was conducted using both non-mutagenized (parent) worms and worms immediately after five generations of mutagen treatment. This assay was performed as an initial phenotypic screen prior to downstream fitness and genomic analyses. Worms from each strain *treatment combination were bleached for age synchronization. The lifetime progeny production of 15 replicates per strain per treatment was monitored by placing a single L4 worm on a 75mm × 13mm Petri plate, transferring it to a fresh plate every 24 hours. After 96 hours, the worm was discarded, and the progeny produced over the four days were sexed and counted. Any worms that died during the transfer process were excluded from the analysis. Male frequency was calculated for each assay plate individually, and treatment means ± SE were calculated across plates.

### Outcrossing frequency assay

Outcrossing frequency was assessed using paired matings (one L4 hermaphrodite + one adult male). For each strain and mutagen treatment, the lifetime progeny production of 15 independent mating plates was monitored, while 30 independent mating plates were assayed for the non-mutagenized parental control. The larger sample size for the non-mutagenized parental control was chosen to provide a high-confidence estimate of baseline male and outcrossing frequencies for each strain, which serve as reference points for classifying outcrossing events in the mutagenized treatments. Each mating pair was placed on a 75 mm × 13 mm Petri plate and transferred to a fresh plate every 24 hours. After 96 hours, both the male and hermaphrodite were discarded, and progeny produced over the four days were allowed to develop to adulthood. Mating plates were pooled across lines sharing the same treatment history.

Because cross-fertilization in *C. elegans* produces approximately 50% male progeny, whereas selfing hermaphrodites produce few or no males, male frequency provides a reliable indicator of whether outcrossing occurred during the assay window ^33^. The goal of this assay was to determine whether successful mating occurred at least once on a plate, rather than to quantify mating efficiency. Plates were therefore classified as either “outcrossing” or “selfing-dominated” using strain-specific criteria informed by baseline male production.

Based on our own baseline measurements, spontaneous male production in the N2 and AB1 strains was consistently below 1% both before and after mutagenesis. For these strains, plates were visually screened for males rather than exhaustively counted. Plates exhibiting male frequencies clearly exceeding the expected spontaneous range (1–5%) were classified as outcrossing. In practice, outcrossed plates were readily distinguishable by the presence of numerous males, whereas selfing-dominated plates contained few or no males and required extended inspection to detect even a single male. In contrast, CB4856 naturally exhibits elevated baseline male frequencies (∼15–17%). For this strain, progeny on each plate were counted and sexed, and male frequency was calculated as the proportion of male progeny. To conservatively distinguish true outcrossing events from baseline variation, CB4856 plates were classified as outcrossing only when male frequency reached or exceeded 25%, a threshold approximately one standard deviation above baseline male frequency observed in this strain. To verify that our classification was not sensitive to the chosen cutoff, we repeated the analysis using alternative thresholds (20% and 30% male frequency) for CB4856. These sensitivity analyses produced qualitatively identical estimates of outcrossing frequency across treatments. Outcrossing frequency was calculated as the proportion of plates meeting these criteria within each strain and treatment.

### Relative fitness assay

After five generations of mutagenesis, worms were frozen at -80°C and later recovered for relative fitness assay at sixth generation-representing the fitness immediately after five generations of mutagenesis and again after three generations of recovery from mutations using the standard *C. elegans* maintenance protocol at 20 °C. During recovery, worms were maintained using the standard worm maintenance protocol. For the relative fitness assay, one early-stage L1 worm from each focal strain (N2, CB4856, AB1) was paired with one L1 worm from the non-mutagenized common competitor strain ST2 expressing green fluorescent protein (GFP). Both strains were co-cultured on NGM plates with *E. coli* OP50 for eight days (∼2 generations) at 20 °C. Washed worms from replicate plates (11 per generation/mutant/strain combination) were collected in M9 on a slide, imaged with and without a GFP filter on a fluorescent microscope, and counted using a Python-based code specific for worm morphology (available on the Github). Plates with zero worms of either type were excluded from analysis, assuming that the worm did not survive to lay eggs. Images captured without the GFP filter counted all worms present on the plate-both the focal strain and the common competitor strain. Conversely, images captured with the GFP filter only revealed the green, fluorescent competitor worms, allowing for estimation of their abundance relative to the total worm population. Relative fitness was quantified from focal and competitor counts using a ratio-based metric as described in the Statistical Analysis section.

Fitness was compared across non-mutagenized, post-mutagenized, and recovered lineages using automated image analysis (Supplementary Material Table 2). To validate the accuracy of the image analysis pipeline, we randomly selected a subset of samples and manually counted individuals. This confirmed that the automated method produced reliable counts, with no observed false positives or false negatives. We found that three generations of recovery from mutagenesis were sufficient for each strain to recover pre-mutagenesis fitness levels for both the mutagens (Figure 1C).

### DNA extraction and Sequencing

To assess the genomic impact of mutagenesis, high molecular weight DNA was extracted from worms (after 3 generations of recovery) using the Cytiva kit (detailed procedure available in the Supplementary Materials). Before starting out the DNA extraction we washed a minimum of ∼10,000 worms with M9 buffer from the plate into a tube and kept the worms rocking overnight to remove associated microbiome. Pooled PacBio long-read sequencing was performed for parental worms of all three strains (N2, CB4856, AB1) and one population per strain per mutagen. Additionally, Illumina short-read sequencing was conducted for each replicate of each treatment (control, EMS, formaldehyde) for all strains. This strategy enabled comprehensive analysis of genomic variants induced by mutagenesis and their phenotypic consequences in subsequent generations.

For the comprehensive genomic analysis, we sequenced a total of 36 samples. Using PacBio long-read sequencing, with an average depth of coverage of 94.52x, we sequenced nine samples: the three parental strains (N2, CB4856, AB1) as controls, and six mutagenized replicates i.e., one replicate per strain treated with each mutagen. Additionally, we performed Illumina short-read sequencing on 27 treatment replicates, achieving an average depth of coverage of 72.11x. These 27 samples included the three parental strains, 12 EMS-treated replicates, and 12 formaldehyde-treated replicates, with four replicates per treatment for each strain. Short-read data were used to identify SNPs, whereas both long- and short-read data were integrated to generate a high-confidence SV dataset.

The male frequency and outcrossing rate assays were conducted on worms derived from the same batch that underwent five generations of mutagenesis. In contrast, the worms used for the relative fitness assay and DNA extraction came from a different set, although they were mutagenized in the same manner as the first batch.

### Population simulations

To quantitatively examine the influence of outcrossing on SV purging and retention we implemented our theoretical model in population simulations using the software SLiM 4.3^64^. Our model described a genomic system where mutations occur as SVs, either insertions, deletions or inversions. Each individual in a population of size *K* carried *n* structural mutations. We constructed dioecious populations containing equal numbers of males and females and self-fertile populations with a proportion of individuals reproducing through outcrossing and a proportion reproducing through selfing. Each population started at a specified size *K* and evolved through overlapping generations with reproduction and fitness differences creating variable population sizes. Each individual had a diploid chromosome 10,000 nucleotides in length. Recombination occurred along the chromosomes with uniform probability at a specified rate *r*. After reproduction SV mutations occurred at a rate µ and were removed at a rate ν, for example by back mutation via gene conversion ^65^.

We did not explicitly model the physical size of mutations but assumed that mutations had discrete phenotypic effect sizes that cumulatively influenced fitness. The probability of a SV mutation occurring at any location was uniform across the chromosome and each SV mutation conferred a small deleterious phenotypic effect drawn from Γ(k, θ) ^39–41^(Supplementary Figure 1). Individual SVs interacted to decrease fitness faster-than-linearly and the fitness landscape had a negative curvature or synergistic epistasis (Supplementary Figure 3). The fitness of an individual carrying n SVs was 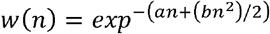 where *a* and *b* were parameters determining the curvature of the fitness landscape ^36^. Our simulated populations evolved for 100,000 generations with the parameters *k* = 1, 0 = 2, *µ* = 0.01, *v* = 0.001, *a* = 0.001 and *b* = 0.0006. We evolved populations under a range of evolutionary scenarios including recombination rates *r*=0.5-1×10^-5^, *K*=100 to 5,000 and levels of self-fertility ranging from 0 to 100%. (Supplementary Figure 2). We also experimented with the curvature of the landscape, modeling a linear fitness landscape as *w*(*n*) = *exp*^*-an*^ with *a* = 0.00635 and a positive or antagonistic curvature where fitness impacts attenuate as they accumulate as 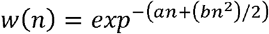 with *a* = 0.001 and *b* = 0.045 (Supplementary Figure 3).

## Data analysis

### Male frequency assay

Male frequency was calculated as the ratio of male progeny to the total progeny laid by each worm, and this proportion was used as the unit of analysis. We assessed the normality of the data using the Shapiro-Wilk test and by visualizing its distribution with a histogram. As expected, the distribution was right-skewed, with a high proportion of zero values, indicating zero-inflation. Given the bounded nature (0 to 1) of the data and the excess zero values, we used a zero-inflated Beta regression for analysis. Since this model requires proportion values strictly between 0 and 1, we applied an epsilon adjustment by adding a small constant to all zero values to make the data more appropriate for the model. We performed the analysis using the glmmTMB package in R ^66^, implementing the following model:

Male Frequency∼Strain× Mutagen

In this model Strain (N2, AB1, CB4856) and Mutagen treatment (EMS, Formaldehyde and non-mutagenized parents were included as fixed factors and an interaction term was added to check if the effect of strain on male frequency varied by mutagen identity. This model accounts for both (1) the continuous variation in male frequency and (2) the excess zero values separately, making it a better fit than standard regression models. Further we did Tukey’s HSD comparison (Abdi & Williams, 2010) on estimated marginal means using emmeans package in R ^68^. For visualization purposes, figures display arithmetic means ± 1 standard error calculated directly from individual assay plates, rather than model-estimated marginal means.

### Relative fitness assay

To quantify the relative fitness of the *C. elegans* strains N2, CB4856, and AB1, each plate was washed with M9 buffer solution to prepare the worms for imaging. Two aliquots from the washed plate were then placed onto a microscope slide and covered with a coverslip. We performed imaging to count the GFP-expressing competitor strain and the non-GFP-expressing focal strain worms within these aliquots. The allelic frequency of the non-fluorescent allele was calculated as the square root of proportion of non-GFP-expressing worms to the total worm population. For statistical analysis, two-way ANOVAs were conducted separately for each mutagen to assess the effects of strain and treatment duration on relative fitness. Data normality was examined using the Shapiro-Wilk test and the residual distribution showed slight variation from normality (Shapiro-Wilk test: W = 0.9868, *p* < 0.01), indicating variation from normality. However, visual inspection of the residuals with histogram suggested an approximately normal distribution. Since ANOVA is robust to minor deviations, we proceeded with ANOVA followed by pairwise comparisons using Tukey’s HSD test. The following linear model was applied:

anova(lm(RF ∼ Strain*Generation, data = a)).

Following ANOVA, we performed pairwise comparisons using Tukey’s Honest Significant Difference (HSD) test to assess differences in relative fitness across strains and generations. We conducted separate Tukey HSD tests within each strain to compare relative fitness before mutagenesis, after five generations of mutagenesis, and after three generations of recovery.

### Bioinformatic analysis

De novo mutations were defined as variants detected in recovered populations that were absent from the corresponding non-mutagenized parental populations when compared with the reference genome. The bioinformatic analysis for identifying *de novo* mutations following PacBio sequencing involved a comprehensive workflow to process and analyse long-read data, distinguishing mutations that arose in the worms after mutagenesis from those existing in parental strains. Initially, raw PacBio reads were aligned and indexed to reference genomes using minimap2 ^69^ and bcftools ^70^; for the N2 and CB4856 strains, NCBI reference genomes were used, while the AB1 strain employed the recently annotated genome of its descendant strain, SX3368^71^, to account for genomic divergences within the lineage. Subsequent variant calling was performed using Sniffles ^72^ for SVs, such as insertions, deletions, and inversions, capitalizing on the accuracy offered by long-read data.

For the specific identification of de novo mutations induced by mutagenesis, a comparative analysis using bcftools contrasted the variant profiles of generation 8 worms (post three generations of recovery) with their control parents, filtering out pre-existing genetic variations. All parental and mutagenized samples were processed using identical library preparation, sequencing, and bioinformatics pipelines to ensure comparability in variant detection. Further quality control and data filtering ensured only high-quality SNPs and SVs (quality score ≥ 20, depth ≥ 10) were retained. We then used bedtools to provide insights into their potential impacts on exonic regions and regulatory elements.

Additionally, similar bioinformatic steps were conducted with Illumina sequencing data. Alignment and indexing were again performed on reference genomes, followed by SNP and SV calling using Freebayes ^73^and Manta ^74^, respectively. We initially called variants in pooled-population mode (FreeBayes ‘pooled’) to account for minor alleles in the combined worm samples. However, when intersecting these SNP calls with structural variants, we observed inflated counts due to multi-allelic sites being represented multiple times. To avoid this over-counting artifact, we generated a separate VCF using non-pooled calling parameters and used these calls for all subsequent intersection and annotation analyses. Enhanced SV detection was achieved with VGtoolkit ^75^ through graph-based variant calling, which was then integrated with SVs identified by Sniffles 2.0 using Survivor 1.0.3^76^, creating a comprehensive view of the genomic alterations induced by the mutagens.

Long-read sequencing was performed on the three non-mutagenized parent strains and one replicate per strain for each mutagen treatment. Short-read sequencing covered all replicates across all strains. For structural variation analysis, we merged variants identified by both long-read and short-read callers using Survivor 1.0.3. For SNP analysis, we relied solely on variants identified by short-read callers. To isolate mutagen-induced effects, the ancestral (pre-treatment) genome was used as the reference baseline for all de novo variant calling. Parallel non-mutagenized controls were not sequenced because spontaneous SVs (SVs) in *C. elegans* occur at extremely low frequencies (approximately 0.003–0.03 events per genome per generation;^42,47^). Given the three-generation duration of our experiment and the detection limits of pooled long-read sequencing, the expected contribution of spontaneous SVs is negligible. Accordingly, the SVs identified here predominantly reflect high-confidence, mutagen-induced genomic changes

To test whether SNPs were enriched within SV regions, the expected number of SNPs within SV intervals was calculated as: Expected = Total SNPs × (SV base pairs / Genome base pairs). Observed SNP counts within SV intervals were then compared with expectation using one-sided binomial tests for each strain–treatment combination.

### Annotation of Transposable Elements and Intersection with Structural Variants

Transposon annotation is facilitated in complete and contiguous sequences, thus annotation of TEs was done in the N2, AB, and CB reference genomes rather than directly from the raw sequence data. We used the TransposonUltimate reasonaTE pipeline for TE annotation in each of the reference genomes. Briefly, TEUltimate reasonaTE wraps 13 previously published TE annotation tools, which include: RepeatMasker v4.1.1, RepeatModeler v2.0.1, SINE-Scan, SINE-Finder, LTRPred, LTRHarvest, TIRvish, MUSTv2, MITE-Tracker, MITEFinderII, HelitronScanner, TransposonPSI for transposon discovery. While some functions are optional, all subprograms were run with default parameters in this study. The resulting annotations are filtered, merged, and clustered using CD-HIT v4.8.1 ^77^ and BLASTN v2.10.1 ^78^. The final annotations from FinalAnnotations_Transposons.gff3 in the final Results folder were used for downstream analysis for each of the three focal strains.

Evidence of transposon movement was assumed from structural variants which overlapped with transposon annotations. The TEUltimatedeTEct pipeline^34^ Click or tap here to enter text.was used for intersection of transposon and structural variant annotations. De novo VCF structural variant files, as generated above, were intersected with transposon annotations and reported as an event if overlapping by at least 10% and lengths were similar by at least 50%. Finally, any annotation smaller than 50 base pairs, or larger than 1% of the genome was discarded. Plots were generated from the transpositionEvents.gff3 file in RStudio with R v4.3.2 (R Core Team, 2023).

## Literature Cited

1. Lynch, M. et al. The Mutational Meltdown in Asexual Populations. We Are Very Grateful To. Journal of Heredity vol. 84 (1993).

2. Lynch, M. et al. Perspective: Spontaneous deleterious mutation. Evolution (N. Y). 53, 645–663 (1999).

3. Robert, F. & Pelletier, J. Exploring the Impact of Single-Nucleotide Polymorphisms on Translation. Frontiers in Genetics vol. 9 Preprint at 10.3389/fgene.2018.00507 (2018).

4. Feuk, L., Carson, A. R. & Scherer, S. W. Structural variation in the human genome. Nature Reviews Genetics vol. 7 85–97 Preprint at 10.1038/nrg1767 (2006).

5. Collins, R. L. et al. A structural variation reference for medical and population genetics. Nature 581, 444–451 (2020).

6. Mahmoud, M. et al. Structural variant calling: The long and the short of it. Genome Biology vol. 20 Preprint at 10.1186/s13059-019-1828-7 (2019).

7. Ho, S. S., Urban, A. E. & Mills, R. E. Structural variation in the sequencing era. Nature Reviews Genetics vol. 21 171–189 Preprint at 10.1038/s41576-019-0180-9 (2020).

8. Glémin, S. How are deleterious mutations purged? Drift versus nonrandom mating. Evolution (N. Y). 57, 2678–2687 (2003).

9. Whitlock, M. C. & Agrawal, A. F. Purging the genome with sexual selection: Reducing mutation load through selection on males. Evolution vol. 63 569–582 Preprint at 10.1111/j.1558-5646.2008.00558.x (2009).

10. García-Dorado, A. Understanding and predicting the fitness decline of shrunk populations: Inbreeding, purging, mutation, and standard selection. Genetics 190, 1461–1476 (2012).

11. Byers, D. L. & Waller, D. M. Do Plant Populations Purge Their Genetic Load? Effects of Population Size and Mating History on Inbreeding Depression. Source: Annual Review of Ecology and Systematics vol. 30 https://about.jstor.org/terms (1999).

12. Szövényi, P. et al. Efficient purging of deleteriousmutations in plants with haploid selfing. Genome Biol. Evol. 6, 1238–1252 (2014).

13. MacPherson, B., Scott, R. & Gras, R. Sex and recombination purge the genome of deleterious alleles: An Individual Based Modeling Approach. Ecological Complexity 45, (2021).

14. Sianta, S. A., Peischl, S., Moeller, D. A. & Brandvain, Y. The efficacy of selection may increase or decrease with selfing depending upon the recombination environment. Evolution (N. Y). 77, 394–408 (2023).

15. Hartfield, M. Evolutionary genetic consequences of facultative sex and outcrossing. J. Evol. Biol. 29, 5–22 (2016).

16. Hoffmann, A. A. & Rieseberg, L. H. Revisiting the impact of inversions in evolution: From population genetic markers to drivers of adaptive shifts and speciation? Annual Review of Ecology, Evolution, and Systematics vol. 39 21–42 Preprint at 10.1146/annurev.ecolsys.39.110707.173532 (2008).

17. Kirkpatrick, M. & Barton, N. Chromosome inversions, local adaptation and speciation. Genetics 173, 419–434 (2006).

18. Kazazian, H. H. Mobile Elements: Drivers of Genome Evolution. https://www.science.org.

19. Feschotte, C. & Pritham, E. J. DNA Transposons and the Evolution of Eukaryotic Genomes.

20. Belyayev, A. Bursts of transposable elements as an evolutionary driving force. Journal of Evolutionary Biology vol. 27 2573–2584 Preprint at 10.1111/jeb.12513 (2014).

21. Wright, S. & Finnegan, D. Genome Evolution: Sex and the Transposable Element. Current Biology vol. 11 (2001).

22. Boutin, T. S., Le Rouzic, A. & Capy, P. How does selfing affect the dynamics of selfish transposable elements? Mob. DNA 3, (2012).

23. Anderson, J. L., Morran, L. T. & Phillips, P. C. Outcrossing and the maintenance of males within C. elegans populations. in Journal of Heredity vol. 101 (2010).

24. Wegewitz, V., Schulenburg, H. & Streit, A. Experimental insight into the proximate causes of male persistence variation among two strains of the androdioecious Caenorhabditis elegans (Nematoda). BMC Ecol. 8, (2008).

25. Manoel, D., Carvalho, S., Phillips, P. C. & Teotónio, H. Selection against males in Caenorhabditis elegans under two mutational treatments. Proceedings of the Royal Society B: Biological Sciences 274, 417–424 (2007).

26. Stewart, A. D. & Phillips, P. C. Selection and Maintenance of Androdioecy in Caenorhabditis Elegans.

27. Gray, J. C. & Cutter, A. D. Mainstreaming Caenorhabditis elegans in experimental evolution. Proceedings of the Royal Society B: Biological Sciences vol. 281 Preprint at 10.1098/rspb.2013.3055 (2014).

28. Manoel, D., Carvalho, S., Phillips, P. C. & Teotónio, H. Selection against males in Caenorhabditis elegans under two mutational treatments. Proceedings of the Royal Society B: Biological Sciences 274, 417–424 (2007).

29. Cutter, A. D. Mutation and the experimental evolution of outcrossing in Caenorhabditis elegans. J. Evol. Biol. 18, 27–34 (2005).

30. Chelo, I. M. et al. Partial selfing can reduce genetic loads while maintaining diversity during experimental evolution. G3: Genes, Genomes, Genetics 9, 2811–2821 (2019).

31. Teterina, A. A. et al. Genomic diversity landscapes in outcrossing and selfing Caenorhabditis nematodes. PLoS Genet. 19, (2023).

32. Kutscher, L. M. & Shaham, S. Forward and reverse mutagenesis in C. elegans. WormBooklZ: the online review of C. elegans biology 1–26 Preprint at 10.1895/wormbook.1.167.1 (2014).

33. Yin, D. & Haag, E. S. Evolution of sex ratio through gene loss. Proc. Natl. Acad. Sci. U. S. A. 116, 12919–12924 (2019).

34. Riehl, K., Riccio, C., Miska, E. A. & Hemberg, M. TransposonUltimate: software for transposon classification, annotation and detection. Nucleic Acids Res. 50, E64 (2022).

35. Kondrashov, A. S. Selection against harmful mutations in large sexual and asexual populations. Genet. Res. 40, 325–332 (1982).

36. Charlesworth, B. OPTIMIZATION MODELS, QUANTITATIVE GENETICS, AND MUTATION. Evolution (N. Y). 44, 520–538 (1990).

37. Charlesworth, B. Mutation-selection balance and the evolutionary advantage of sex and recombination. Genet. Res. 55, 199–221 (1990).

38. Haller, B. C. & Messer, P. W. SLiM 4: Multispecies Eco-Evolutionary Modeling. American Naturalist 201, E127–E139 (2023).

39. Eyre-Walker, A. & Keightley, P. D. The distribution of fitness effects of new mutations. Nature Reviews Genetics vol. 8 610–618 Preprint at 10.1038/nrg2146 (2007).

40. Keightley, P. D. & Eyre-Walker, A. Joint inference of the distribution of fitness effects of deleterious mutations and population demography based on nucleotide polymorphism frequencies. Genetics 177, 2251–2261 (2007).

41. Racimo, F. & Schraiber, J. G. Approximation to the Distribution of Fitness Effects across Functional Categories in Human Segregating Polymorphisms. PLoS Genet. 10, (2014).

42. Konrad, A. et al. Mitochondrial mutation rate, spectrum and heteroplasmy in Caenorhabditis elegans spontaneous mutation accumulation lines of differing population size. Mol. Biol. Evol. 34, 1319–1334 (2017).

43. Estes, S., Phillips, P. C., Denver, D. R., Thomas, W. K. & Lynch, M. Mutation Accumulation in Populations of Varying Size: The Distribution of Mutational Effects for Fitness Correlates in Caenorhabditis Elegans.

44. Ciccia, A. & Elledge, S. J. The DNA Damage Response: Making It Safe to Play with Knives. Molecular Cell vol. 40 179–204 Preprint at 10.1016/j.molcel.2010.09.019 (2010).

45. Limpose, K. L., Corbett, A. H. & Doetsch, P. W. BERing the burden of damage: Pathway crosstalk and posttranslational modification of base excision repair proteins regulate DNA damage management. DNA Repair vol. 56 51–64 Preprint at 10.1016/j.dnarep.2017.06.007 (2017).

46. Hillier, L. D. W. et al. Comparison of C. elegans and C. briggsae genome sequences reveals extensive conservation of chromosome organization and synteny. PLoS Biol. 5, 1603–1616 (2007).

47. Saxena, A. S. & Baer, C. F. High rate of mutation and efficient removal by selection of structural variants from natural populations of Caenorhabditis elegans. Preprint at 10.1101/2025.03.22.644739 (2025).

48. Kim, C. et al. Long-read sequencing reveals intra-species tolerance of substantial structural variations and new subtelomere formation in C. elegans. Genome Res. 29, 1023–1035 (2019).

49. Bush, Z. D. et al. Transposable elements and heterochromatic regions are enriched for structural variation and sequence divergence in the genome of wild-type Caenorhabditis elegans. G3: Genes, Genomes, Genetics 15, (2025).

50. Thompson, O. A. et al. Remarkably divergent regions punctuate the genome assembly of the Caenorhabditis elegans hawaiian strain CB4856. Genetics 200, 975–989 (2015).

51. Brenner, S. THE GENETICS OF CAENORHABDZTZS ELEGANS.

52. Andersen, E. C. et al. Chromosome-scale selective sweeps shape Caenorhabditis elegans genomic diversity. Nat. Genet. 44, 285–290 (2012).

53. Ristow, M. & Schmeisser, K. Mitohormesis: Promoting health and lifespan by increased levels of reactive oxygen species (ROS). Dose-Response 12, 288–341 (2014).

54. Cypser, J. R. & Johnson, T. E. Multiple Stressors in Caenorhabditis Elegans Induce Stress Hormesis and Extended Longevity. Journal of Gerontology vol. 57 https://academic.oup.com/biomedgerontology/article/57/3/B109/550368 (2002).

55. Negi, P., Rai, A. N. & Suprasanna, P. Moving through the stressed genome: Emerging regulatory roles for transposons in plant stress response. Frontiers in Plant Science vol. 7 Preprint at 10.3389/fpls.2016.01448 (2016).

56. Wright, S. I. & Schoen, D. J. Transposon Dynamics and the Breeding System. Genetica vol. 107 (1999).

57. Dolgin, E. S. & Charlesworth, B. The fate of transposable elements in asexual populations. Genetics 174, 817–827 (2006).

58. Emmons, S. W. & Yesner, L. High-Frequency Excision of Transposable Element Tel in the Nematode Caenorhabditis Elegans Is Limited to Somatic Cells. Cell vol. 36 (1964).

59. Mori, I., Benian, G. M., Moermant, D. G. & Waterston, R. H. Transposable Element Tcl of Caenorhabditis Elegans Recognizes Specific Target Sequences for Integration (Site Specificity/Transposon/Unc-22/Nematode). Proc. Natl. Acad. Sci. USA vol. 85 (1988).

60. Bao, W., Jurka, M. G., Kapitonov, V. V. & Jurka, J. New superfamilies of eukaryotic DNA tyransposons and their internal divisions. Mol. Biol. Evol. 26, 983–993 (2009).

61. Laricchia, K. M., Zdraljevic, S., Cook, D. E. & Andersen, E. C. Natural Variation in the Distribution and Abundance of Transposable Elements Across the Caenorhabditis elegans Species. Mol. Biol. Evol. 34, 2187–2202 (2017).

62. Sterken, M. G., Snoek, L. B., Kammenga, J. E. & Andersen, E. C. The laboratory domestication of Caenorhabditis elegans. Trends in Genetics vol. 31 224–231 Preprint at 10.1016/j.tig.2015.02.009 (2015).

63. Anderson, J. L., Morran, L. T. & Phillips, P. C. Outcrossing and the maintenance of males within C. elegans populations. in Journal of Heredity vol. 101 (2010).

64. Haller, B. C. & Messer, P. W. SLiM 4: Multispecies Eco-Evolutionary Modeling. American Naturalist 201, E127–E139 (2023).

65. Katju, V., LaBeau, E. M., Lipinski, K. J. & Bergthorsson, U. Sex change by gene conversion in a Caenorhabditis elegans fog-2 mutant. Genetics 180, 669–672 (2008).

66. Bolker, B. Getting Started with the GlmmTMB Package. (2022).

67. Abdi, H. & Williams, L. J. Encyclopedia of Research Design. http://www.utd.edu/~herve.

68. Searle, S. R., Speed, F. M. & Milliken, G. A. Population marginal means in the linear model: An alternative to least squares means. American Statistician 34, 216–221 (1980).

69. Li, H. Minimap2: Pairwise alignment for nucleotide sequences. Bioinformatics 34, 3094–3100 (2018).

70. Danecek, P. et al. Twelve years of SAMtools and BCFtools. Gigascience 10, (2021).

71. Riccio C. Riccio, C., 2022. Duplication is a prominent mechanism of recent gene birth in Caenorhabditis elegans (Doctoral dissertation). (2022).

72. Smolka, M. et al. Detection of mosaic and population-level structural variants with Sniffles2. Nat. Biotechnol. https://doi.org/10.1038/s41587-023-02024-y (2024) doi:10.1038/s41587-023-02024-y.

73. Garrison, E. & Marth, G. Haplotype-based variant detection from short-read sequencing. http://arxiv.org/abs/1207.3907 (2012).

74. Chen, X. et al. Manta: Rapid detection of structural variants and indels for germline and cancer sequencing applications. Bioinformatics 32, 1220–1222 (2016).

75. Hickey, G. et al. Genotyping structural variants in pangenome graphs using the vg toolkit. Genome Biol. 21, (2020).

76. Jeffares, D. C. et al. Transient structural variations have strong effects on quantitative traits and reproductive isolation in fission yeast. Nat. Commun. 8, (2017).

77. Fu, L., Niu, B., Zhu, Z., Wu, S. & Li, W. CD-HIT: Accelerated for clustering the next-generation sequencing data. Bioinformatics 28, 3150–3152 (2012).

78. Camacho, C. et al. BLAST+: Architecture and applications. BMC Bioinformatics 10, (2009).

79. Riehl, K., Riccio, C., Miska, E. A. & Hemberg, M. TransposonUltimate: software for transposon classification, annotation and detection. Nucleic Acids Res. 50, E64 (2022).

