## Supplementary figures tables and protocols for "Strain-specific structural variant landscapes shape mutation retention following mutagenesis in *Caenorhabditis elegans*"

**Supplementary Materials**

1. ***Protocols***

**DNA Extraction Protocol – Sera-Xtracta HMW DNA Kit**

Preparation

- Wash the *C. elegans* plates thoroughly with M9 buffer to collect worms.
- Transfer the worm suspension to a tube and rock gently overnight at room temperature to allow removal of surface-associated microbiota.
- Centrifuge the suspension to pellet the worms.
- Carefully discard the supernatant without disturbing the pellet.
- Add fresh M9 buffer to the pellet and repeat the wash to further remove residual bacteria.
- Centrifuge again and discard the supernatant.
- Resuspend the final worm pellet in 200 µL M9 buffer in a microcentrifuge tube.
- Add 0.6 µL of 10% SDS solution to lyse the worms.
- Subject the samples to five freeze–thaw cycles:
- Freeze at –80 °C, then thaw at 25 °C, repeating this process five times to enhance lysis.

.

### Sample Lysis

- 1. Add 20 µL Proteinase K to the bottom of a 2 mL microcentrifuge tube.
- 2. Add 200 µL of worm sample
- 3. Add 200 µL Lysis buffer and vortex for 15 seconds.
- 4. Incubate at 25°C for 30 minutes (no agitation).
- 5. Vortex briefly for 5 seconds post incubation.
- 6. Briefly centrifuge to collect the lysate at the bottom of the tube.

### DNA Binding

- 1. Add 15 µL magnetic bead suspension (vortex before use).
- 2. Add 230 µL Binding buffer and vortex for 5 seconds.
- 3. Incubate at 25°C for 3 minutes at 1400 rpm.
- 4. Briefly centrifuge and place the tube on a magnetic rack for 1 minute.
- 5. Carefully aspirate and discard the supernatant.

### Washing (4 total washes)

- 1. Remove the tube from the magnetic rack.
- 2. Add 700 µL Wash 1 buffer. Vortex or shake to disperse beads.
- 3. Incubate at 25°C for 1 minute at 1400 rpm.
- 4. Centrifuge briefly and place on magnetic rack. Remove supernatant.
- 5. Repeat steps 1–4 with Wash 1 buffer once more.
- 6. Repeat steps 1–4 twice with Wash 2 buffer.

### Drying (Optional but recommended)

- 1. Briefly centrifuge to collect droplets.
- 2. Place on magnetic rack and aspirate any residual wash buffer.
- 3. Air dry the bead pellet for 5 minutes.

### Elution

- 1. Remove from magnet and add 100 µL Elution buffer.
- 2. Vortex to mix and dislodge pellet.
- 3. Incubate at 25°C for 3 minutes at 1400 rpm.
- 4. Place on magnetic rack for 1 minute.
- 5. Transfer the eluted DNA (supernatant) to a clean tube.

### Storage

- Store purified DNA at 4°C for short-term use.
- For long-term storage, aliquot and freeze at -20°C or lower.
- Avoid repeated freeze-thaw cycles.

**Supplementary Table 1: Pairwise contrasts of male frequency between *C. elegans* strains (AB1, CB4856, N2) and treatments (non-mutagenized parental controls, EMS, and formaldehyde), based on estimated marginal means from a zero-inflated beta regression model. Estimates represent differences on the model (logit) scale, with associated standard errors (SE), z-ratios, and Tukey-adjusted *p*-values derived from post hoc comparisons using the *emmeans* framework.**

| **Contrast** | **estimate** | **SE** | **z.ratio** | **p.value** |
| --- | --- | --- | --- | --- |
| AB1 EMS - CB4856 EMS | -0.305 | 0.409 | -0.746 | 0.9981 |
| AB1 EMS - N2 EMS | 0.1547 | 0.395 | 0.392 | 1 |
| AB1 EMS - AB1 Formaldehyde | -0.021 | 0.401 | -0.052 | 1 |
| AB1 EMS - CB4856 Formaldehyde | -0.4269 | 0.427 | -1 | 0.986 |
| AB1 EMS - N2 Formaldehyde | 0.2724 | 0.394 | 0.691 | 0.9989 |
| AB1 EMS - AB1 Parent | 0.0301 | 0.36 | 0.083 | 1 |
| AB1 EMS - CB4856 Parent | -1.4614 | 0.358 | -4.085 | 0.0015 |
| AB1 EMS - N2 Parent | 0.1396 | 0.348 | 0.401 | 1 |
| CB4856 EMS - N2 EMS | 0.4597 | 0.387 | 1.188 | 0.9592 |
| CB4856 EMS - AB1 Formaldehyde | 0.284 | 0.394 | 0.722 | 0.9985 |
| CB4856 EMS - CB4856 Formaldehyde | -0.1219 | 0.42 | -0.291 | 1 |
| CB4856 EMS - N2 Formaldehyde | 0.5774 | 0.387 | 1.493 | 0.8591 |
| CB4856 EMS - AB1 Parent | 0.3351 | 0.352 | 0.952 | 0.9899 |
| CB4856 EMS - CB4856 Parent | -1.1564 | 0.348 | -3.319 | 0.0254 |
| CB4856 EMS - N2 Parent | 0.4446 | 0.339 | 1.31 | 0.9287 |
| N2 EMS - AB1 Formaldehyde | -0.1757 | 0.379 | -0.464 | 0.9999 |
| N2 EMS - CB4856 Formaldehyde | -0.5816 | 0.406 | -1.433 | 0.8852 |
| N2 EMS - N2 Formaldehyde | 0.1177 | 0.371 | 0.317 | 1 |
| N2 EMS - AB1 Parent | -0.1246 | 0.335 | -0.372 | 1 |
| N2 EMS - CB4856 Parent | -1.6161 | 0.333 | -4.859 | <.0001 |
| N2 EMS - N2 Parent | -0.0151 | 0.322 | -0.047 | 1 |
| AB1 Formaldehyde - CB4856 Formaldehyde | -0.4059 | 0.412 | -0.985 | 0.9873 |
| AB1 Formaldehyde - N2 Formaldehyde | 0.2934 | 0.378 | 0.776 | 0.9975 |
| AB1 Formaldehyde - AB1 Parent | 0.0511 | 0.343 | 0.149 | 1 |
| AB1 Formaldehyde - CB4856 Parent | -1.4404 | 0.34 | -4.236 | 0.0008 |
| AB1 Formaldehyde - N2 Parent | 0.1606 | 0.33 | 0.487 | 0.9999 |
| CB4856 Formaldehyde - N2 Formaldehyde | 0.6993 | 0.406 | 1.724 | 0.7317 |
| CB4856 Formaldehyde - AB1 Parent | 0.457 | 0.373 | 1.226 | 0.9509 |
| CB4856 Formaldehyde - CB4856 Parent | -1.0345 | 0.369 | -2.804 | 0.1141 |
| CB4856 Formaldehyde - N2 Parent | 0.5665 | 0.361 | 1.57 | 0.8212 |
| N2 Formaldehyde - AB1 Parent | -0.2423 | 0.335 | -0.724 | 0.9985 |
| N2 Formaldehyde - CB4856 Parent | -1.7338 | 0.333 | -5.213 | <.0001 |
| N2 Formaldehyde - N2 Parent | -0.1328 | 0.321 | -0.413 | 1 |
| AB1 Parent - CB4856 Parent | -1.4915 | 0.291 | -5.125 | <.0001 |
| AB1 Parent - N2 Parent | 0.1095 | 0.279 | 0.393 | 1 |
| CB4856 Parent - N2 Parent | 1.601 | 0.276 | 5.803 | <.0001 |

**Supplementary Table 2. Pairwise comparisons of relative fitness across non-mutagenized, mutagenized, and recovered populations for each strain and mutagen treatment. Comparisons were performed using Tukey’s HSD post hoc tests following the ANOVA framework described in the Statistical Analysis section. Values are differences in mean relative fitness between groups (diff), with corresponding confidence bounds (lwr, upr) and Tukey-adjusted p-values (p adj).**

| **Strain** | **Mutagen** | **Comparison** | **Difference (diff)** | **Lower Bound (lwr)** | **Upper Bound (upr)** | **P-Value (p adj)** |
| --- | --- | --- | --- | --- | --- | --- |
| N2 | Formaldehyde | Non-mutagenised - Mutagenised | -0.1207 | -0.2298 | -0.0116 | 0.02639 |
| N2 | Formaldehyde | Recovered - Mutagenised | -0.1166 | -0.2118 | -0.0213 | 0.01235 |
| N2 | Formaldehyde | Recovered - Non-mutagenised | 0.0041 | -0.1012 | 0.1095 | 0.99518 |
| N2 | EMS | Non-mutagenised - Mutagenised | -0.0554 | -0.1657 | 0.0549 | 0.45756 |
| N2 | EMS | Recovered - Mutagenised | -0.0541 | -0.1554 | 0.0472 | 0.41321 |
| N2 | EMS | Recovered - Non-mutagenised | 0.0012 | -0.1098 | 0.1123 | 0.99961 |
| CB4856 | Formaldehyde | Non-mutagenised - Mutagenised | 0.0283 | -0.0841 | 0.1406 | 0.82078 |
| CB4856 | Formaldehyde | Recovered - Mutagenised | 0.0282 | -0.0915 | 0.1479 | 0.84065 |
| CB4856 | Formaldehyde | Recovered - Non-mutagenised | -4.00E-05 | -0.1076 | 0.1075 | 0.99999 |
| CB4856 | EMS | Non-mutagenised - Mutagenised | 0.0081 | -0.1011 | 0.1173 | 0.98291 |
| CB4856 | EMS | Recovered - Mutagenised | -0.0736 | -0.1903 | 0.0432 | 0.29531 |
| CB4856 | EMS | Recovered - Non-mutagenised | -0.0817 | -0.1897 | 0.0264 | 0.17516 |
| AB1 | Formaldehyde | Non-mutagenised - Mutagenised | -0.0014 | -0.1279 | 0.1251 | 0.9996 |
| AB1 | Formaldehyde | Recovered - Mutagenised | 0.015 | -0.0961 | 0.1261 | 0.94408 |
| AB1 | Formaldehyde | Recovered - Non-mutagenised | 0.0164 | -0.0986 | 0.1314 | 0.93766 |
| AB1 | EMS | Non-mutagenised - Mutagenised | 0.0931 | -0.0369 | 0.223 | 0.20632 |
| AB1 | EMS | Recovered - Mutagenised | 0.1276 | 0.0155 | 0.2397 | 0.02183 |
| AB1 | EMS | Recovered - Non-mutagenised | 0.0345 | -0.0817 | 0.1507 | 0.75683 |

**Supplementary material Table 3. Structural variant classes detected after mutagenesis.**

| **Strain** | **Treatment** | **Intra INS/DEL/DUP/INV (mean ± SD)** | **Interchromosomal TRA (mean ± SD)** | **TRA as % of total (mean)** |
| --- | --- | --- | --- | --- |
| AB1 | EMS | 120.00 ± 0.00 | 1.00 ± 0.00 | 0.83 |
| AB1 | Formal | 95.00 ± 0.00 | 3.00 ± 0.00 | 3.06 |
| CB4856 | EMS | 917.25 ± 0.50 | 29.00 ± 0.00 | 3.06 |
| CB4856 | Formal | 816.50 ± 0.58 | 37.00 ± 0.00 | 4.34 |
| N2 | EMS | 519.00 ± 0.00 | 4.00 ± 0.00 | 0.76 |
| N2 | Formal | 542.00 ± 0.00 | 4.00 ± 0.00 | 0.73 |

**Supplementary Table 4. Aggregated enrichment of SNPs within structural variant regions across strains and mutagen treatments.**

| **Strain** | **Treatment** | **Total SNPs** | **Observed SNPs in SV** | **SV Fraction** | **Expected SNPs in SV** | **Enrichment (Obs/Exp)** | **p-value (binomial test)** |
| --- | --- | --- | --- | --- | --- | --- | --- |
| AB1 | EMS | 28,124 | 3,377 | 0.0406 | 1,140.05 | 2.96× | < 1 × 10⁻³⁰⁰ |
| AB1 | Formaldehyde | 28,211 | 1,795 | 0.0248 | 700.53 | 2.56× | 6.33 × 10⁻²⁷¹ |
| CB4856 | EMS | 40,954 | 33,769 | 0.4312 | 17,667.73 | 1.91× | < 1 × 10⁻³⁰⁰ |
| CB4856 | Formaldehyde | 40,130 | 30,952 | 0.4066 | 16,316.40 | 1.90× | < 1 × 10⁻³⁰⁰ |
| N2 | EMS | 27,333 | 1,947 | 0.0272 | 744.46 | 2.62× | 6.15 × 10⁻³⁰⁶ |
| N2 | Formaldehyde | 30,724 | 2,090 | 0.0140 | 429.55 | 4.87× | < 1 × 10⁻³⁰⁰ |

**Supplementary Table 5*: Strain-specific distribution of structural variant breakpoint microhomology lengths***

| **Strain** | **0 bp (NHEJ)** | **1–2 bp (NHEJ/MMEJ)** | **3–10 bp (MMEJ)** | **11–50 bp (SSA)** | **>50 bp (homology mediated repair)** |
| --- | --- | --- | --- | --- | --- |
| **AB1** | 0.234 | 0.091 | 0.083 | 0.285 | 0.307 |
| **CB4856** | 0.202 | 0.096 | 0.029 | 0.142 | 0.531 |
| **N2** | 0.153 | 0.086 | 0.043 | 0.108 | 0.610 |

**Supplementary Table 6*: Very few TEs were involved in transposition events***

| **STRAIN** | **TREATMENT** | **TOTAL TE COUNT** | **Transposition Events** | **PERCENT of TEs in Transposition Events** |
| --- | --- | --- | --- | --- |
| **N2** | EMS | 15533 | 11 | 0.071 |
| **N2** | Formal | 15533 | 13 | 0.084 |
| **AB1** | EMS | 23375 | 10 | 0.043 |
| **AB1** | Formal | 23375 | 13 | 0.056 |
| **CB4856** | EMS | 15155 | 42 | 0.277 |
| **CB4856** | Formal | 15155 | 37 | 0.244 |

**Supplementary Table 7*: SV mutations above 10kb account for a small percent of all SV mutations***

| **STRAIN** | **TREATMENT** | **SV>10kb COUNT** | **TOTAL SVs** | **PERCENT of SVs>10kb** |
| --- | --- | --- | --- | --- |
| **AB1** | Formal | 2 | 140 | 1.4 |
| **AB1** | EMS | 2 | 164 | 1.2 |
| **N2** | Formal | 9 | 634 | 1.4 |
| **N2** | EMS | 9 | 607 | 1.5 |
| **CB4856** | Formal | 17 | 1218 | 1.4 |
| **CB4856** | EMS | 18 | 1307 | 1.4 |

Supplementary Material Table 8: Nucleotides affected by structural variations for each strain

| **Strain** | **Mutagen** | **Insertion** | **Deletion** | **Duplication** | **Inversion** |
| --- | --- | --- | --- | --- | --- |
| N2 -1 | Formaldehyde | 471238 | 74044 | 44318 | 1289368 |
| N2 -2 | Formaldehyde | 471238 | 74044 | 44318 | 1289368 |
| N2 -3 | Formaldehyde | 471238 | 74044 | 44318 | 1289368 |
| N2 -4 | Formaldehyde | 471238 | 74044 | 44318 | 1289368 |
| N2 -1 | EMS | 482307 | 56936 | 36357 | 2642474 |
| N2 -2 | EMS | 482307 | 56936 | 36357 | 2642474 |
| N2 -3 | EMS | 482307 | 56936 | 36357 | 2642474 |
| N2 -4 | EMS | 482307 | 56936 | 36357 | 2642474 |
| CB4856-1 | Formaldehyde | 960440 | 118305 | 1169792 | 49702760 |
| CB4856-2 | Formaldehyde | 960182 | 118422 | 1169792 | 49702760 |
| CB4856-3 | Formaldehyde | 960237 | 118305 | 1169792 | 49702760 |
| CB4856-4 | Formaldehyde | 960409 | 118422 | 1169792 | 49702760 |
| CB4856-1 | EMS | 1011363 | 152738 | 21170220 | 32829363 |
| CB4856-2 | EMS | 1011427 | 152738 | 21170220 | 32829363 |
| CB4856-3 | EMS | 1011363 | 152738 | 21170220 | 32829363 |
| CB4856-4 | EMS | 1011363 | 152738 | 21170220 | 32829363 |
| AB-1 | Formaldehyde | 35468 | 46363 | 0 | 2539375 |
| AB-2 | Formaldehyde | 35468 | 46363 | 0 | 2539375 |
| AB-3 | Formaldehyde | 35468 | 46363 | 0 | 2539375 |
| AB-4 | Formaldehyde | 35468 | 46363 | 0 | 2539375 |
| AB-1 | EMS | 28269 | 22975 | 4640 | 4181603 |
| AB-2 | EMS | 28269 | 22975 | 4640 | 4181603 |
| AB-3 | EMS | 28269 | 22975 | 4640 | 4181603 |
| AB-4 | EMS | 28269 | 22975 | 4640 | 4181603 |

**Supplementary Table 9*: SV mutations above 10kb***

| **STRAIN** | **TREATMENT** | **SV** | **SV SIZE** | **CHR** | **POSITION** |
| --- | --- | --- | --- | --- | --- |
| **AB1** | Formal | DEL | -17180 | V | 20076945 |
| **AB1** | Formal | INV | 2539375 | V | 20059968 |
| **AB1** | EMS | INV | 1558731 | IV | 2379651 |
| **AB1** | EMS | INV | 2539369 | V | 20059962 |
| **N2** | EMS | DEL | -13014 | V | 6094837 |
| **N2** | EMS | INS | 10852 | I | 3990048 |
| **N2** | EMS | INS | 11122 | V | 12509981 |
| **N2** | EMS | DUP | 16801 | III | 1267035 |
| **N2** | EMS | INS | 17826 | II | 3994672 |
| **N2** | EMS | INV | 33856 | IV | 3170028 |
| **N2** | EMS | INV | 39175 | X | 256072 |
| **N2** | EMS | INV | 78335 | II | 7124208 |
| **N2** | EMS | INV | 2488588 | X | 15796614 |
| **N2** | Formal | DEL | -13014 | V | 6094837 |
| **N2** | Formal | DEL | -12521 | IV | 16121858 |
| **N2** | Formal | INS | 10847 | I | 3990048 |
| **N2** | Formal | INS | 16121 | V | 12510218 |
| **N2** | Formal | DUP | 16801 | III | 1267035 |
| **N2** | Formal | INV | 43288 | IV | 6733414 |
| **N2** | Formal | INV | 70146 | V | 15106170 |
| **N2** | Formal | INV | 78335 | II | 7124208 |
| **N2** | Formal | INV | 1095182 | IV | 9538168 |
| **CB4856** | EMS | DEL | -12538 | V | 18975224 |
| **CB4856** | EMS | INS | 10445 | V | 17027656 |
| **CB4856** | EMS | INS | 10461 | V | 18942522 |
| **CB4856** | EMS | INS | 11418 | X | 7175802 |
| **CB4856** | EMS | INS | 11920 | IV | 14689338 |
| **CB4856** | EMS | INS | 13261 | II | 410527 |
| **CB4856** | EMS | DUP | 13397 | V | 8183016 |
| **CB4856** | EMS | INS | 13466 | X | 4878846 |
| **CB4856** | EMS | INV | 17259 | V | 17128283 |
| **CB4856** | EMS | INV | 31306 | IV | 1303580 |
| **CB4856** | EMS | INV | 711398 | II | 1732522 |
| **CB4856** | EMS | INV | 6287897 | II | 7253397 |
| **CB4856** | EMS | INV | 7100396 | V | 19165121 |
| **CB4856** | EMS | INV | 8605254 | I | 13063839 |
| **CB4856** | EMS | DUP | 8823231 | I | 10475255 |
| **CB4856** | EMS | INV | 10054305 | II | 12621065 |
| **CB4856** | EMS | DUP | 12307181 | IV | 14046466 |
| **CB4856** | Formal | DEL | -12538 | V | 18975224 |
| **CB4856** | Formal | DEL | -11928 | X | 16291 |
| **CB4856** | Formal | INS | 10413 | V | 18942522 |
| **CB4856** | Formal | INS | 11418 | X | 7175802 |
| **CB4856** | Formal | INS | 12932 | X | 4878736 |
| **CB4856** | Formal | DUP | 13397 | V | 8183016 |
| **CB4856** | Formal | INV | 17259 | V | 17128283 |
| **CB4856** | Formal | INS | 18068 | II | 3866976 |
| **CB4856** | Formal | INS | 18245 | II | 3867133 |
| **CB4856** | Formal | INV | 104174 | II | 1016307 |
| **CB4856** | Formal | INV | 244930 | X | 1611520 |
| **CB4856** | Formal | INV | 711398 | II | 1732522 |
| **CB4856** | Formal | DUP | 1127283 | V | 17626148 |
| **CB4856** | Formal | INV | 6287897 | II | 7253397 |
| **CB4856** | Formal | INV | 7100396 | V | 19165121 |
| **CB4856** | Formal | INV | 10054305 | II | 12621065 |
| **CB4856** | Formal | INV | 11239626 | III | 12145539 |
| **CB4856** | Formal | INV | 13914551 | V | 16195689 |

##### Supplementary Table 10. Median and mean sizes of mutagen-induced insertions and deletions across strains: *Summary statistics are calculated from replicate-level structural variant (SV) calls.*

| **Strain** | **Mutagen** | **SV type** | **Mean SV size (bp)** | **Mean median SV size (bp)** | **Mean SV count per replicate** |
| --- | --- | --- | --- | --- | --- |
| AB1 | EMS | Deletion | 298.38 | 118.00 | 77.0 |
| AB1 | EMS | Insertion | 724.85 | 287.00 | 39.0 |
| AB1 | Formaldehyde | Deletion | 833.34 | 140.50 | 56.0 |
| AB1 | Formaldehyde | Insertion | 933.37 | 239.50 | 38.0 |
| N2 | EMS | Deletion | 382.12 | 184.00 | 149.0 |
| N2 | EMS | Insertion | 1351.00 | 435.00 | 357.0 |
| N2 | Formaldehyde | Deletion | 435.55 | 184.00 | 170.0 |
| N2 | Formaldehyde | Insertion | 1319.99 | 381.00 | 357.0 |
| CB4856 | EMS | Deletion | 526.68 | 173.00 | 290.0 |
| CB4856 | EMS | Insertion | 1707.69 | 693.50 | 592.2 |
| CB4856 | Formaldehyde | Deletion | 496.28 | 173.00 | 238.5 |
| CB4856 | Formaldehyde | Insertion | 1791.64 | 766.50 | 536.0 |

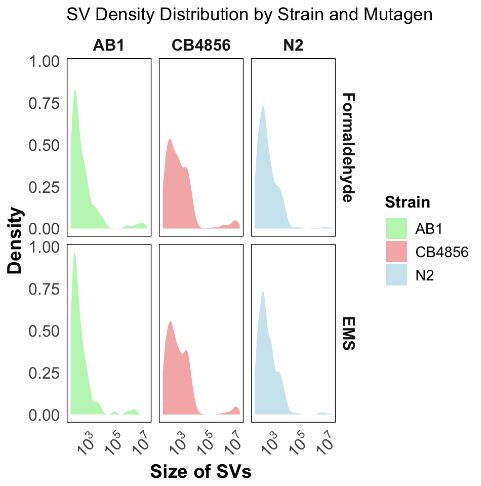

Supplementary Figure 1: Size distribution of structural variants (SVs) across *C. elegans* strains and mutagens: Density distribution of SVs sizes for all the three strains for both the mutagens. x- axis Size of SVs, Y-axis: density of SVs.

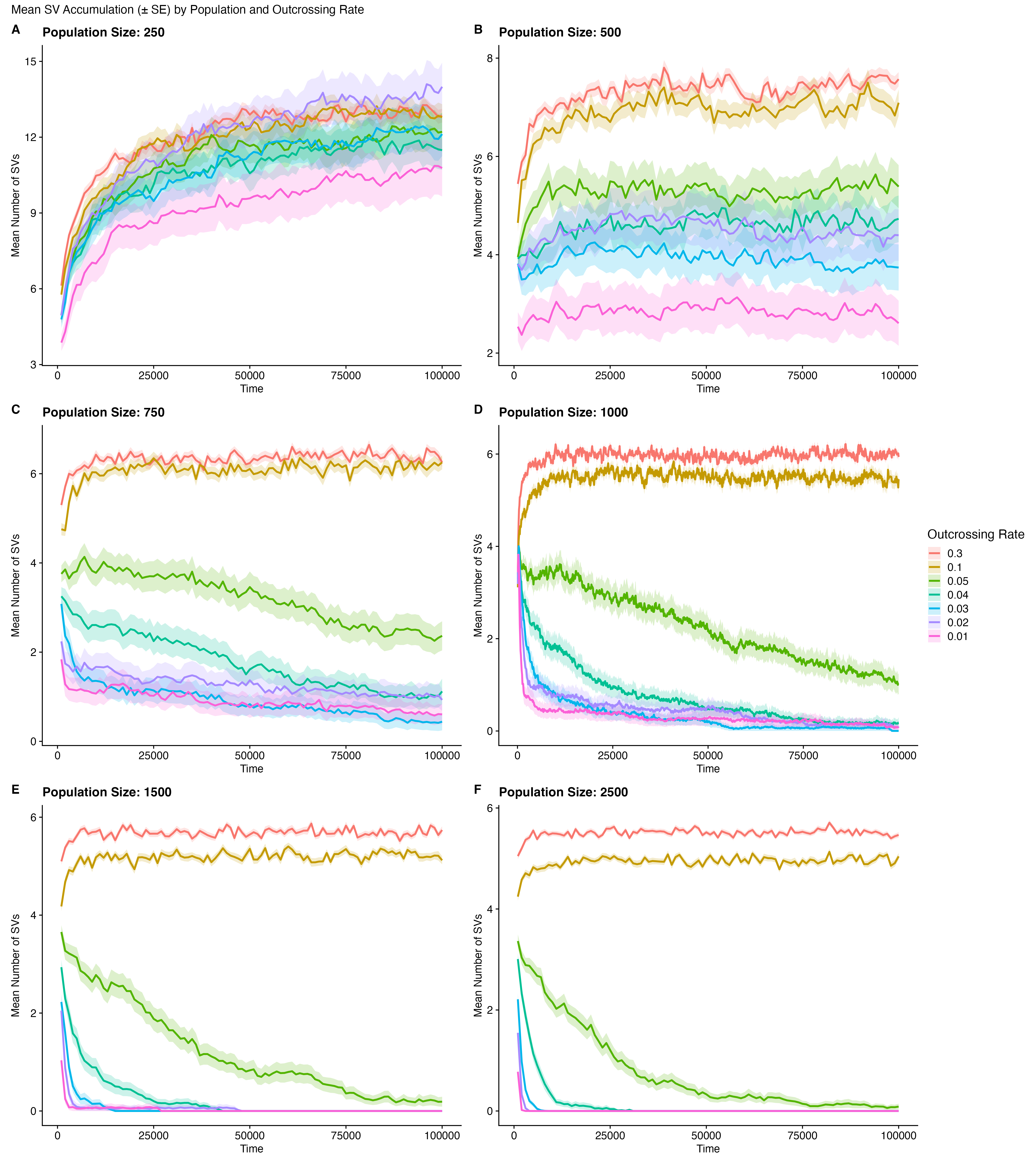

**Supplementary Figure 2:** Simulation of structural variant (SV) accumulation across population sizes and outcrossing rates under a gamma-distributed distribution of fitness effects: Mean SV number (± SE) is shown over time for simulated populations with varying sizes (panels A–F: 250, 500, 750, 1000, 1500, and 2500 individuals) and outcrossing rates (color-coded). Mutations were modeled as having small deleterious effects drawn from a gamma distribution. Across smaller population sizes and higher outcrossing rates, SVs accumulated and were maintained at higher levels, whereas larger populations and reduced outcrossing facilitated more efficient purging. Shaded regions indicate standard error across replicate simulations.

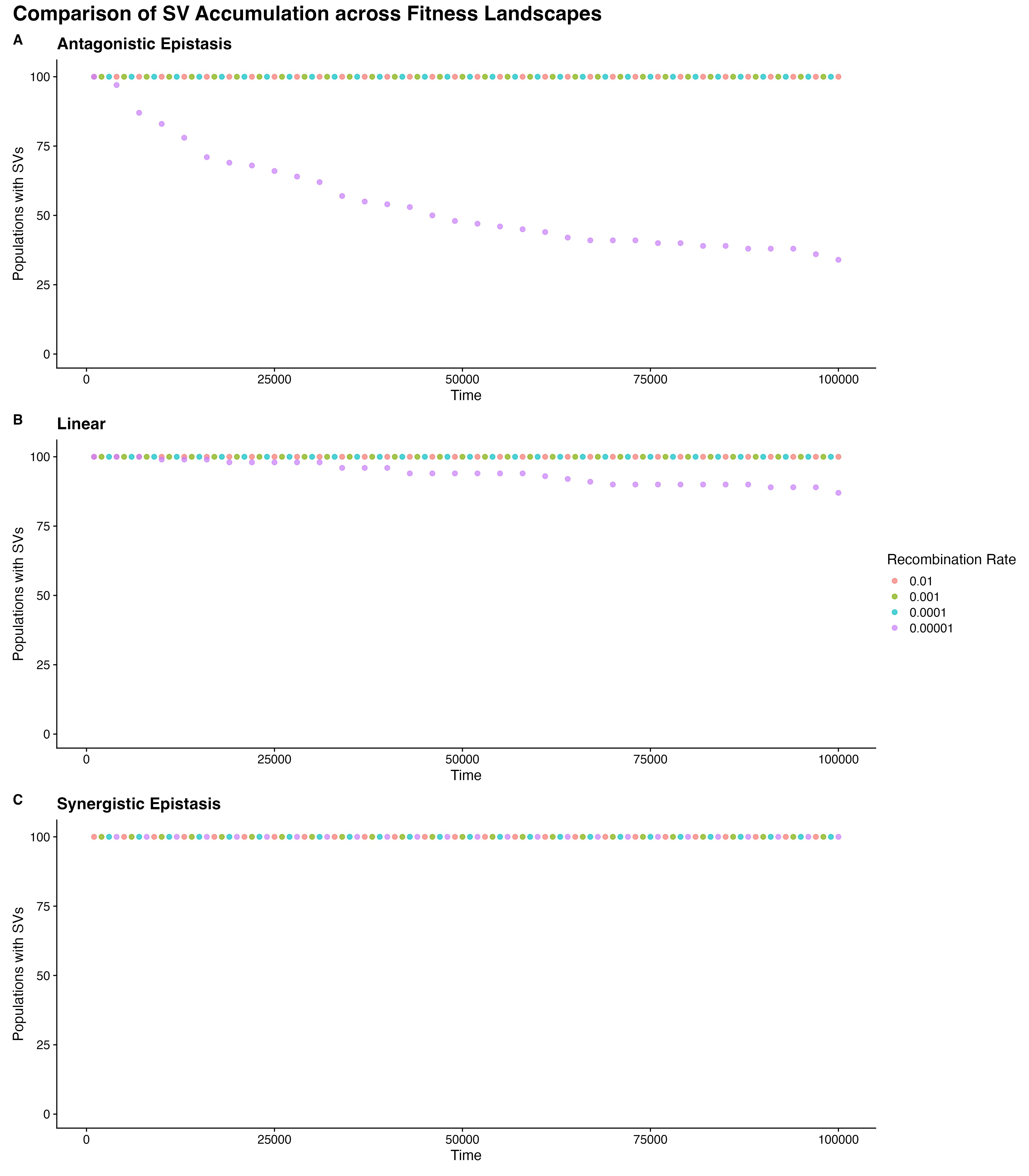

**Supplementary Figure 3:**Effect of fitness landscape shape on structural variant (SV) persistence in simulated populations.: The proportion of simulated populations retaining SVs over time is shown under three fitness landscape models: (A) antagonistic epistasis, (B) linear effects, and (C) synergistic epistasis (negative curvature). Colors indicate recombination rates. Under antagonistic and linear fitness landscapes, populations largely retained SVs across all recombination rates, whereas under synergistic epistasis, SV persistence remained high across conditions, indicating that negative curvature does not promote efficient purging in these simulations.
